# Uncertainty mapping and probabilistic tractography using Simulation-based Inference in diffusion MRI: A comparison with classical Bayes

**DOI:** 10.1101/2024.11.19.624267

**Authors:** J.P. Manzano-Patron, Michael Deistler, Cornelius Schröder, Theodore Kypraios, Pedro J. Gonçalves, Jakob H. Macke, Stamatios N. Sotiropoulos

## Abstract

Simulation-Based Inference (SBI) has recently emerged as a powerful framework for Bayesian inference: Neural networks are trained on simulations from a forward model, and learn to rapidly estimate posterior distributions. We here present an SBI framework for parametric spherical deconvolution of diffusion MRI data of the brain. We demonstrate its utility for estimating white matter fibre orientations, mapping uncertainty of voxel-based estimates and performing probabilistic tractography by spatially propagating fibre orientation uncertainty. We conduct an extensive comparison against established Bayesian methods based on Markov-Chain Monte-Carlo (MCMC) and find that: a) in-silico training can lead to calibrated SBI networks with accurate parameter estimates and uncertainty mapping for both single- and multi-shell diffusion MRI, b) SBI allows amortised inference of the posterior distribution of model parameters given unseen observations, which is orders of magnitude faster than MCMC, c) SBI-based tractography yields reconstructions that have a high level of agreement with their MCMC-based counterparts, equal to or higher than scan-rescan reproducibility of estimates. We further demonstrate how SBI design considerations (such as dealing with noise, defining priors and handling model selection) can affect performance, allowing us to identify optimal practices. Taken together, our results show that SBI provides a powerful alternative to classical Bayesian inference approaches for fast and accurate model estimation and uncertainty mapping in MRI.

## 1. Introduction

Identifying the parameters of a model that best represent a set of observations is an omnipresent task in science. For Magnetic Resonance Imaging (MRI) in particular, this is a key task, as on many occasions biophysical quantities of interest are indirectly inferred from measurements. In diffusion MRI (dMRI) of the brain, measurements of the diffusion scatter pattern of water molecules within tissues can be mapped through models to neuronal tissue microstructure and brain connectivity [1, 2, 3].

Deterministic approaches, from non-linear optimisation to machine learning methods (see [4] for a review), can be used to fit such models to data, providing *point estimates* of model parameters. However, these estimates are affected by different sources of uncertainty, from noise in the observations (aleatoric uncertainty), to preprocessing errors, modelling assumptions, and incomplete sampling (epistemic uncertainty). Mapping uncertainty provides a principled manner for assessing confidence in the estimates [5], separating noise and signal, identifying covariance structures in the parameters and aiding experimental design [6]. In dMRI, uncertainty in fibre orientations has been the basis for probabilistic tractography [7, 8, 9, 10], which estimates spatial distributions of white matter bundles.

Classical approaches for mapping uncertainty in dMRI include bootstrapping and Bayesian inference. Bootstrapping [11] involves iteratively fitting the same model to randomly resampled (or perturbed) versions of the original dataset. This provides an empirical distribution for each model parameter, whose width denotes the respective estimation uncertainty [12, 13, 14, 15], but can be sensitive to how resampling is performed [16, 17]. On the other hand, Bayesian approaches offer a principled framework for estimating the posterior distribution of the model parameters given the data, providing measures of confidence and covariance in the estimates. For instance, Variational Bayes [18, 19, 20], or Markov Chain Monte Carlo (MCMC) sampling [21, 22, 23] provide flexibility in modelling both signal and noise components, and make it possible to explicitly incorporate prior knowledge. These approaches have a number of limitations: 1) they require the likelihood of the model to be evaluated analytically, which is not always available or it is too expensive to compute, 2) although great speed-ups have been achieved (e.g. utilising GPUs [24] and/or improving the algorithmic design [25, 26]), they can be highly iterative in nature, which can result in substantial computational complexity and limited scalability, 3) convergence to sampling the correct posterior can be a challenge.

Simulation-based inference (SBI) [27] has been developed to tackle such limitations in Bayesian inference. The basic idea is that artificial neural networks are trained on simulations from a forward model, in a way that allows them to learn Bayesian posterior distributions. Importantly, this approach can *amortise* inference: after initial training, performing inference on unseen data only requires a single forward pass through the trained neural network, without the need for costly iterative sampling and likelihood evaluations [28, 29]. This amortisation property is particularly effective in scenarios where the same problem needs to be solved many times, as in the voxel-wise inference done in microstructural models of dMRI.

In this work, we demonstrate how SBI can be used for mapping orientations and uncertainty in dMRI and, for the first time, all the way up to probabilistic tractography. We comprehensively compare SBI with a classical MCMC-based and heavily-used implementation (FSL-BedpostX) [7, 30, 31], with respect to both voxel-wise fibre orientation uncertainty estimates and whole-brain white matter tract reconstructions.

### 1.1. Related work

In Bayesian inference, the posterior distribution *p*(**Ω**|*Y*_*obs*_) of model parameters **Ω** given the measured data *Y*_*obs*_ is proportional to the product of the prior *p*(**Ω**) and the likelihood function *p*(*Y*_*obs*_|**Ω**), the conditional distribution of observations given a model. For problems where the likelihood is intractable or expensive to compute [32, 33], Approximate Bayesian Computation (ABC) or Likelihood-Free Inference (LFI) approaches have been proposed [34, 35]. In ABC, the posterior is approximated by simulating large amounts of data *Y*_*sim*_ from a forward stochastic simulator *f*_*Y*_ (**Ω**) (synthetic likelihood), with parameters **Ω** randomly drawn from the defined priors *p*(**Ω**). Samples for which a distance *ρ*(*Y*_*sim*_, *Y*_*obs*_) is below a pre-defined threshold are accepted as posterior samples (otherwise rejected). SBI recasts this as a density estimation problem, where neural networks are used to learn a mapping between observations and the posterior distribution [35, 36]. This avoids inefficient sampling-rejection schemes and allows problems with high-dimensional observations to be tackled. Such networks, known as Neural Density Estimators (NDEs) [37], have enabled inference in a purely data-driven fashion on different target (conditional) densities, such as the posterior distribution of model parameters (Neural Posterior Estimation or NPE) [38, 39, 40, 41], a surrogate likelihood (Neural Likelihood Estimation) [42, 43], or variants based on denoising diffusion models [44, 45, 46, 47].

A few recent studies have explored inferring parameters from dMRI models using such approaches, demonstrating how SBI can assist in fitting models that are typically challenging to invert. Jallais et al. [48] fitted the SANDI model of grey matter [49] to direction-averaged dMRI signal, using the width of the SBI-estimated posteriors to identify regions where the model presented indeterminacies or biases. To reduce dimensionality and avoid indeterminacies, they employed rotationally invariant dMRI summary statistics to train SBI networks on synthetic data (relying on stacked Masked Autoencoders (MADE)), demonstrating accurate parameter estimation given appropriate q-space sampling. A generalisation of this work was presented in [50], proposing a toolbox applicable to a broader range of models with examples of fitting to direction-averaged dMRI signals. In Eggl and De Santis [51], the authors used the mean of the SBI posterior to demonstrate that it can provide robust point-estimates with fewer observations than deterministic non-linear fitting. In another recent work, Karimi et al. [52] presented a likelihood-free framework to obtain the full posterior of parameters in a multi-tensor model. This employed a two-step strategy: a convolutional neural network (CNN) for model selection (classifying dMRI signals into single or multi-fibre), followed by Mixture Density Networks (MDNs) with Gaussian kernels trained for inference against simulations obtained from the multi-tensor model. This work used a spherical harmonics (SH) representation of the signals improving generalisability of inference across different acquisition schemes. An extension of this work for the standard model of white matter [53] was presented in [54], demonstrating the ability to obtain fiber-specific microstructural features.

However, despite the great potential revealed for SBI from these studies, a comprehensive comparison to classical Bayesian approaches, which have been used in dMRI for more than two decades, is still missing. In particular, Bayesian frameworks for mapping orientation uncertainty and performing uncertainty-based probabilistic tractography [21, 7, 30, 24] have been collectively used in thousands of studies before [10]. Therefore, they naturally provide a reference against which SBI approaches can be calibrated, optimised and validated.

### 1.2. Contributions

In this study we close this gap, expanding on our previous preliminary work [55] and complementing recent work by others [48, 50, 52, 54]. We devise an SBI framework for mapping fibre orientation uncertainty from diffusion MRI data using parametric spherical deconvolution models. We subsequently use the SBI-derived voxel-wise posterior distributions to perform probabilistic tractography for the first time. Given the inherent difficulty in validating uncertainty estimates, we comprehensively evaluate the accuracy and precision of the estimates against an established MCMC-based classical inference approach, crucially using the same models and priors, as implemented in FSL-BedpostX [7, 30, 31]. We evaluate SBI both for voxel-wise and whole-brain level inference, exploring how estimated uncertainty is spatially propagated.

We find that in-silico training can lead to SBI networks that yield accurate mean parameter estimates and uncertainty for unseen observations in both single- and multi-shell dMRI acquisitions, while offering orders of magnitude computational speed-ups for inference. We also show that SBI-based probabilistic tractography for a representative set of more than 40 white matter bundles exhibits a high level of agreement with MCMC-based counterparts; and that this level of agreement is within the range of individual scan-rescan reproducibility of estimates. We further identify optimal SBI design features (such as how to account for noise, how to restrict and constraint priors and how to deal with model selection) that improve training, increase performance and similarity to MCMC estimates. Our findings augment recent studies suggesting that SBI provides new and exciting opportunities for amortised inference in MRI models.

## 2. Methods

### 2.1. NPE theory and network architecture

In classical Bayesian inference (like MCMC, see Fig. 1), given a signal and noise model, a likelihood function *p*(*Y*_*obs*_|**Ω**) can be defined. Any prior knowledge of the parameters **Ω** can be embedded in the prior *p*(**Ω**) and the two can be used to estimate the posterior distribution of the parameters given the observed data, i.e. *p*(**Ω**|*Y*_*obs*_) ∼ *p*(*Y*_*obs*_|**Ω**)*p*(**Ω**). This typically necessitates highly iterative processes and potentially expensive likelihood calculations. Furthermore, parameter estimation given a dataset starts from scratch for any new observations *Y*_*obs*_ (e.g. data for different voxels).

**Figure 1:**
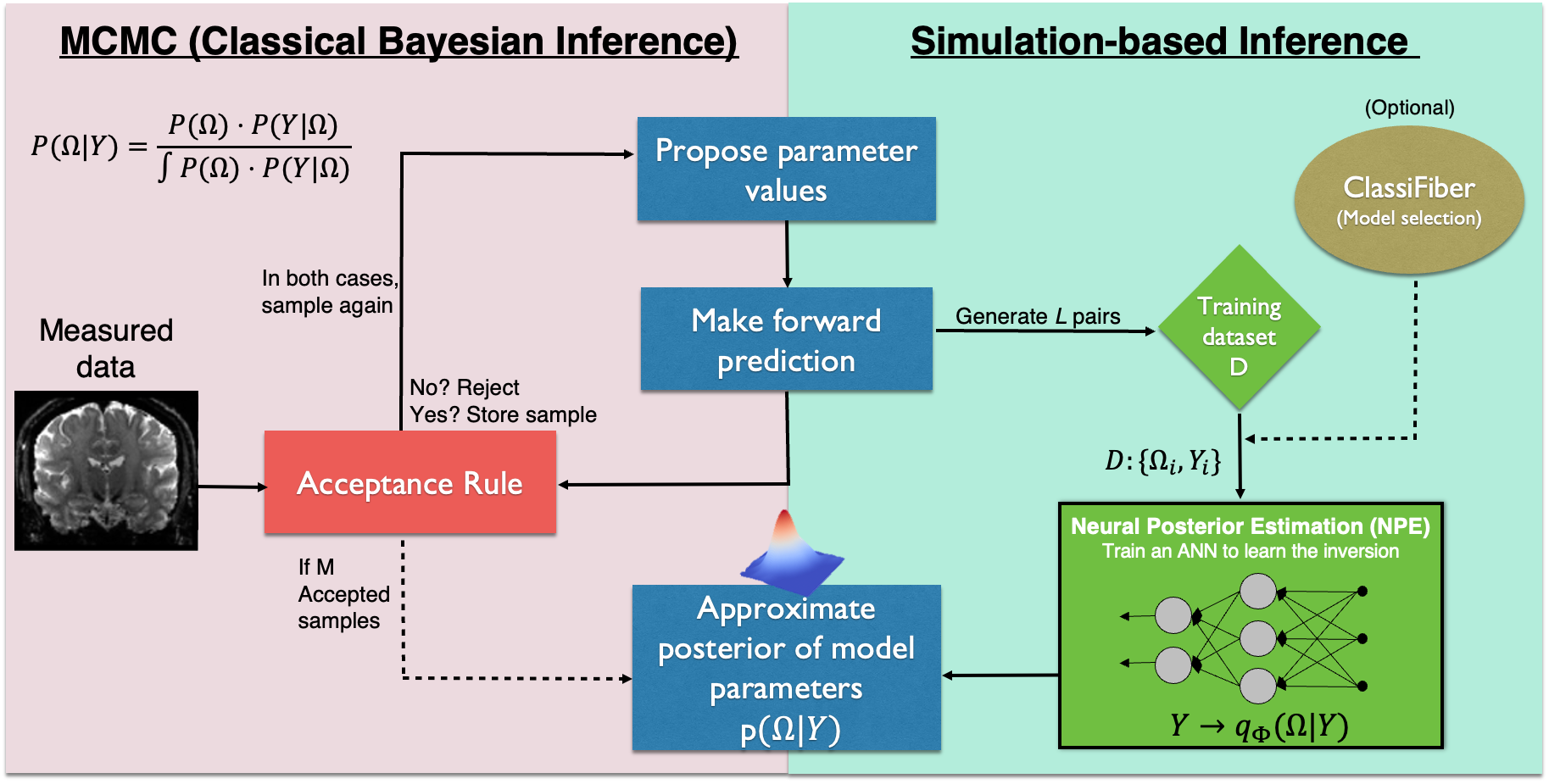
Schematic overview of Markov-Chain Monte-Carlo (MCMC) (an example of Classical Bayesian Inference) and Simulation-Based Inference (SBI). For NPE, an artificial neural network (ANN) is trained on a set {**Ω**_***i***_, ***Y***_***i***_} obtained from the forward simulator, to learn the latent mapping *q*_Φ_ between observations and posterior samples.

Within SBI, this problem can be addressed using Neural Posterior Estimation (NPE) [38, 39, 41]. In NPE, the mapping between observations and posterior is learned once using a training dataset and can be applied on new unseen data without any expensive calculations, a property known as *amortised inference*. Being able to evaluate the likelihood function is not necessary as long as we have a forward process *f*_*Y*_ (**Ω**) to generate simulated data *Y*. As in all Bayesian inference schemes, we can code prior knowledge into our prior distribution *p*(**Ω**) to define what regions of the parameter space are sampled for such forward predictions (or *simulations*). NPE approximates the target posterior using a parameterised conditional density distribution *q*_Φ_(**Ω**|*Y*) represented by neural networks with parameters Φ. Hence, having a training set of pairs (Ω_*i*_, *Y*_*i*_), a NPE is trained to learn the mapping function *q*_Φ_(**Ω**|*Y*) ≈ *p*(**Ω**|*Y*). After training, we can use *q*_Φ_(**Ω**|*Y*) to obtain samples from the (approximate) posterior *p*(**Ω**|*Y*_*obs*_) with a single forward pass through the network.

More formally, we minimise the expected Kullback-Leibler divergence between true posterior distribution *p*(**Ω**|*Y*) and the approximation *q*_Φ_(**Ω**|*Y*) [38, 56]:

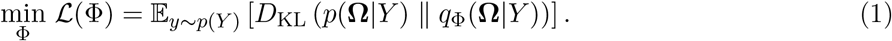

This is equivalent to minimising the negative log-likelihood of the posterior approximation [41, 56]:

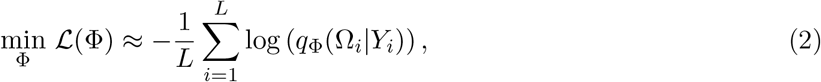

where *L* denotes the number of pair-vectors (**Ω**_*i*_, *Y*_*i*_) used to train the networks, where **Ω** ∼ *p*(**Ω**) and *Y*_*i*_ ∼ *f*_*Y*_ (**Ω**_*i*_). During training, the parameters Φ of the network are optimised to minimise the loss in eq. 2.

Different model choices have been made to parameterise the posterior distribution *q*_Φ_(**Ω**|*Y*) in previous applications to dMRI, such as Mixture Density Networks (MDNs) [52, 54] and Masked Autoencoders (MADE) [48, 50]. In our work, we use Normalising Flows (NFs) as the basis structure for the density estimators given their flexibility and superior performance in previous studies [57, 58, 55]. NFs can produce complex probability distributions by transforming a simple base distribution through a series of invertible transformations *f*_*k*_. A key feature of NFs is their use of differentiable closed-form functions that support efficient computation of both the transformations and their Jacobian determinants, ensuring rapid density estimation and sampling. In this paper, we use Neural Spline Flows (NSF) as the basis function [59], as implemented in the SBI package [60]. NSFs comprise *K* rational-quadratic splines, each parameterised by a neural network.

### 2.2. Forward model f_Y_ (Ω)

In this work, we use variants of the Ball & Sticks model [7, 30] as the forward simulator *f*_*Y*_ (**Ω**). Our goal is to infer the posterior of parameters **Ω** from dMRI data in each voxel, with *Y*_*obs*_ = {*Y*_1,*obs*_, …, *Y*_*j,obs*_} of *j* = 1, …, *J* signals, each corresponding to a different diffusion-sensitising gradient. This model has been heavily used for performing parametric spherical deconvolution and uncertainty-based probabilistic tractography over the past 20 years. An MCMC-based implementation for classical Bayesian inference on this model has been developed by members of our team and is part of FSL [7, 31, 24] and will be used for direct comparisons with SBI.

The forward model is multi-compartment, comprising of a weighted sum of *N* anisotropic compartments (sticks), representing distinct white matter fibre compartments, and an isotropic compartment (ball), representing partial volume and non-axonal contributions to the signal. The voxel-wise dMRI signal attenuation *A* in single-shell data is described as:

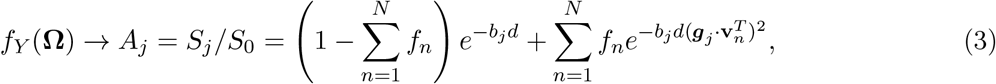

where *S*_*j*_ and *S*_0_ are the signals measured with and without diffusion-sensitising gradients, respectively, with *b*_*j*_ value and orientation ***g***_*j*_ (protocol-defined parameters). The unknown model parameters to estimate are the mean diffusivity *d*, the vector **v**_*n*_ describing the *n*^th^ fibre orientation, represented in spherical coordinates by [*θ*_*n*_, *ϕ*_*n*_] (*θ* is the angle from the positive z-axis and *ϕ* is the angle of the projection on the x-y plane from the positive x-axis) and the volume fraction of the *n*^th^ fibre *f*_*n*_ (with ∑ *f*_*n*_ ≤ 1). The maximum number of fibres *N* per voxel is typically set to *N* = 3, but the appropriate *N* per voxel needs to be determined (model selection challenge).

The above model can be extended for multi-shell data [30] as:

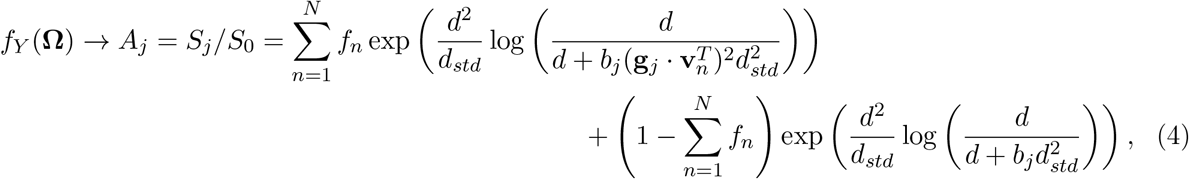

where *d* and *d*_*std*_ denote the mode and spread of a Gamma distribution of diffusivities that can account for non-monoexponential signal decay. In summary, the vector of model parameters when the maximum number of fibres is *n*, 1 ≤ *n* ≤ *N*. is given by **Ω** = {*d, d*_*std*_, {*f*_1_, …, *f*_*n*_, *θ*_1_, …, *θ*_*n*_, *ϕ*_1_, …, *ϕ*_*n*_}}.

### 2.3. SBI design and implementation

We designed and implemented a number of NPE networks for the forward models described in the previous section (setting *N* up to 3), using the Python SBI toolbox (v.0.22) [60]. We set the depth of the NPE to *K* = 10 transformations. For training and inference, we used the attenuation *A*_*j*_ as input for the networks, obtained after data pre-processing (in case of real data), by dividing the diffusion-weighted signal by the mean of the *b* = 0 volumes (without loss of generality, we set *S*_0_=1 for the simulated training sets). Normalised signals generally provide more stable behaviour on neural networks, as the *S*_0_ intensities are not quantitative and can be arbitrarily large or small, depending on the scaling factor applied by the scanner. Depending on model complexity and number of compartments considered, different training sizes were used for different SBI architectures (2M for single-fibre models, 4M for *N* = 2 fibres, and 6M when *N* = 3). Once the networks were trained, we drew *M* = 50 samples from *q*_Φ_(Ω|*Y*_*obs*_) to characterise the posterior given unseen observations, following the MCMC implementation that draws M=50 thinned samples from the posterior [7] (notice that none of the trends in the results changed with more samples). Code for the implemented architectures can be found on https://github.com/orgs/SPMIC-UoN/SBI_dMRI.

We performed extensive comparisons between posterior estimates obtained from SBI and from MCMC (as available in FSL-BedpostX implementation (v.6.0.7) [24]) on both single and multi-shell models. These included comparisons using both synthetic and in-vivo data, at a voxel (e.g., fibre orientations) and whole brain scale (tractography reconstructions). The equations and parameters underlying the MCMC implementation are provided in Supplementary Material for completeness, as well as the metrics we used for assessing accuracy and precision of parameter estimates.

We also investigated how SBI design choices affect accuracy and precision, and agreement with MCMC. In the following subsections we present novel explorations on the training design for handling priors, treating noise and addressing model selection challenges. We finally present the data used and data processing aspects all the way to probabilistic tractography.

#### 2.3.1. Uniform vs restricted priors p(Ω)

Non-informative priors have been used in recent dMRI SBI implementations [48, 50, 52, 54], as they provide the simplest choice for training and offer uniform coverage over the whole parameter space. However, certain combinations of parameters will never be found in real-world cases or would not be feasible to detect due to limitations in the measurement process. Furthermore, certain model parameters have non-linear boundary conditions that cannot be easily defined by parametric prior distributions, requiring ad-hoc reparametrisation that often are not trivial [50]. For instance, the sum of volume fractions in a multi-compartment model should be between 0 and 1. Using non-informative priors and forcing NPE to learn from implausible regions in parameter space could make training highly inefficient, potentially reduce the predictive performance, and increase estimation uncertainty.

We tested two SBI designs, one with uniform priors and another with restricted priors. While the former is straightforward to implement, the latter requires a more advanced design [61, 62]. For the uniform prior implementation we used: *p*(*d*) *∼* 𝒰 (10^−5^, 5 · 10^−3^), *p*(*d*_*std*_) *∼* 𝒰 (0, 5 · 10^−3^),red *p*(*f*_1_) *∼* 𝒰 (0, 1), and *p*(*f*_*n*_) *∼* 𝒰 (0.05, 1). The fibre orientations were sampled in the positive hemisphere (i.e. *θ* ∈ [0, *π/*2], *ϕ* ∈ [0, *π*)) to avoid degenerate solutions due to antipodal symmetry. To ensure uniform sampling on the sphere, the following correction was applied: *θ* = arccos(1 − 2*u*) and *ϕ* = *π* · *u*, where *u* ∼ *U* (0, 1) (see Supplementary Material and Suppl. Fig. S1).

Subsequently, we designed and trained a Restricted Prior distribution that learns feasible regions of the parameter space delimited by (non-linear) constraints on top of the parameter prior, as summarised in Table 1 (soft constraints based on empirical evidence and prior studies on the models considered here ([7, 30]) to avoid degeneracies). To achieve that, we trained a classifier to learn what is the signal stemming from “valid” parameter combinations, i.e. combinations for which restrictions and boundary conditions on individual parameters are respected (see Suppl. Fig. S2). To do so, we 1) sampled each parameter independently from uniform priors *p*(**Ω**) specified above, 2) used these samples to generate forward predictions using *f*_*Y*_ (**Ω**), 3) tagged the cases that did not meet the boundary conditions into a different class of “non-valid”, and 4) trained a classifier accordingly to separate valid vs non-valid cases (see Fig. S2). This classifier was wrapped into a density distribution object to produce samples only from the (“valid”) restricted regions of the parameter space. This trained Restricted Prior object is a joint prior distribution that produces samples for all model parameters at once, in a way that they fulfill all conditions imposed. We trained separate Restricted Priors for the cases of 1, 2 and 3 fibres, and compared the efficiency and performance of the inference against the network using the default uniform priors.

**Table 1:** Parameter constraints and restricted priors in the Ball&Sticks model. These ensure that i) the sum of the volume fractions ranges within [0, 1], ii) there is some ordering in the volume fractions *f*_1_ *> f*_2_ *> f*_3_, iii) detectable crossing angles cannot be lower than a minimum, reflecting limitations in angular contrast/resolution in the data.

|  | Parameter constraints |  |  |
| --- | --- | --- | --- |
|  | 1 Fibre | 2 Fibres | 3 Fibres |
| $f_1$ | $\sim U[0, 1]$ | $\sim U[0.05, 1]$ | |
| $f_2$ | $\sim U[0, 0.001]$ | $\sim U[0.05, 1]$ | |
| $f_3$ | $\sim U[0, 0.001]$ | | $\sim U[0.05, 1]$ |
| $f_{sum}$ | $f_1 \leq 1$ | $(f_1 + f_2) \leq 1$ | $(f_1 + f_2 + f_3) \leq 1$ |
| $f_n$ | None | $f_1 \geq f_2$ | $f_1 \geq f_2 \geq f_3$ |
| $f_n - f_m$ ($m > n$) | None | $> 0.05$ | |
| $\text{angle}(\mathbf{v}_n, \mathbf{v}_m)$ | None | $> 10^\circ$ | $> 30^\circ$ |
| $d_{std}/d$ | $\leq 2$ | | |

#### 2.3.2. Dealing with noise during training

Noisy realisations of the signal were considered during training. For consistency with the MCMC implementation, we assumed zero-mean Gaussian additive noise (i.e. *f*_*Y*_ (**Ω**) → *f*_*Y*_ (**Ω**) + *ε*) with *ε* ∼ 𝒩 (0, *σ*), *σ* = *S*_0_*/SNR* and *SNR* the signal-to-noise ratio of the *b* = 0 signal.

Matching the noise distribution between training and test data improves robustness and accuracy of SBI estimates [63]. However, as such matching cannot be performed a-priori for a generalisable implementation, we evaluated the performance of SBI under two strategies for considering noise during training. For a given set of parameter values **Ω**_***i***_ we explored two strategies: a) Single-level noise: A single noisy realisation *Y*_*i*_ was generated with an SNR value chosen randomly from a prior *p*(*SNR*) ∼ 𝒰 (3, 80), b) Multi-level noise: The same noise-free signal was corrupted with different noise levels chosen from multiple predefined SNR intervals. Here we used 8 SNR intervals: *p*(*SNR*_1_) ∼ 𝒰 (3, 10), *p*(*SNR*_2_) ∼ 𝒰 (10, 20), … *p*(*SNR*_8_) ∼ 𝒰 (70, 80). I.e. we produce 8 noisy realisations of each *Y*_*i*_, each at a different noise level. We kept the total number of training points equal for both strategies. Hence, the former approach provided for training more combinations of parameters sampled from the priors, each with a single noisy realisation; while the latter approach provided (8-times) fewer combinations of parameters for training, but with more examples of noise contamination for each combination.

#### 2.3.3. Model selection using SBI

In the multi-compartment model considered, there is an inherent challenge of deciding how many fibre compartments (i.e. orientations) must be estimated in each voxel. This is a model selection problem that the MCMC implementation addresses using Automatic Relevance Determination (ARD) priors [7] (see Suppl. Material). These are sparsifying improper priors that cannot be sampled from, hence not well suited nor easily adaptable to simulations from the prior, as needed in SBI. Hence, to address the model selection problem with SBI, we have implemented and evaluated two strategies:

A. ClassiFiber + Individual NPEs: Similar to [52, 64], we subdivided the problem into 2 stages: an initial classification step to select how many fibre compartments correspond to the signal of each voxel, and a second step that fits the corresponding model based on the classifier output. For the first step, we designed the *ClassiFiber*, a feed-forward deep neural net that can take the dMRI signal (or a basis signal representation projection) as input and provide the number of fibres as output (with *N* = 1, 2, 3). The neural network is formed by 6 layers (input, 256, 128, 64, 32, output), with ReLu activation functions, 1% dropout, adaptive learning rate, Adam optimisation, and a Cross-Entropy loss function. We generated 6 million parameter combinations (2 million for *N* =1, 2 and 3 crossing-fibres configurations sampled from the restricted priors) and generated signal predictions using the multi-shell model. For the second step, we trained 3 independent NPEs for models with *N* =1, 2, or 3 fibre compartments, respectively, using the basis structure of sec.2.1. Hence, a network e.g. with *N* = 2 compartments was fed with only 2-way crossing patterns during training, while a network with *N* = 1 compartment was not trained with any fibre crossing examples. Note that training data with *N* = 1, as generated using *p*(*f*_1_) ∼𝒰 (0, 1), considered cases where *f*_1_ was zero or very close to zero. Therefore, they implicitly included a set of *N* = 0 cases, where no fibres were present and the signal was dominated by the ball compartment. We refer to this setup as *SBI_ClassiFiber*.
B. Joint NPE: Contrary to the independent NPEs trained above, we trained a network using a training super-set that included examples of 1,2 and 3 crossing-fibre cases (equal number of 1, 2, and 3 fibre cases). In order to keep a fixed parameter vector size, *N* = 3 compartments were set into the model. For cases where *N <* 3 fibre compartments were simulated (i.e. for single fibres and 2-way crossings), the respective volume fractions of the *n*^*th*^ degenerate compartments in the training data were set to *f*_*n*_ ∼ 𝒰 (0, 1*e* − 3), while the values for the orientations (*θ*_*n*_, *ϕ*_*n*_) were randomly sampled from the priors (e.g. when two fibres were simulated, the volume fraction of the third compartment was set to almost zero in the training data). This joint dataset was used to train a single NPE, which we evaluated on whether it can inherently learn the correct number of compartments, without a prior classification as above. Similar to *SBI*_*ClassiFiber, N* = 0 cases were implicitly considered in the *N* = 1 training set, with *f*_1_ ∼ 0.. We refer to this setup as *SBI_joint*.

Note that this joint NPE approach does not directly estimate the number of compartments *N*. Instead, a model with a maximum number of compartments *N*_*max*_ = 3 is fitted and the number of compartments supported by the data is found indirectly by a-posteriori setting a minimum threshold to the estimated volume fractions *f*_*i*_ *> f*_*cutoff*_, *i* = 2, 3. We have used *f*_*cutoff*_ =5%, as similarly used in the MCMC implementation [7].

### 2.4. Datasets, processing and tractography

#### Data

We evaluated the different architectures using both synthetic and in-vivo data. Preprocessed in-vivo dMRI data were obtained from a previous study [65], consisting of 6 re-scans of the same subject, each acquired following a UKBiobank-like multi-shell protocol (2mm isotropic resolution, TR=3s, TE=92ms, MB=3, Siemens Prisma 3T scanner) [66]. Fifty gradient directions at b=1000 s/mm_2_ and another fifty gradient directions at b=2000 s/mm_2_ were obtained (i.e. a total of 100 datapoints per voxel), with a mean SNR of 30 for the b=0 s/mm_2_ image. Diffusion-weighted volumes were normalized by the mean of the *b*_0_ volumes to obtain the signal attenuation. Synthetic data allowed toy examples for testing the inference frameworks under scenarios where the true parameter values were known. We used the attenuation of the signal (with *S*_0_ = 1) from the Ball&Sticks model to generate such synthetic datasets, following the same acquisition protocol (bvals and bvecs) as in the in-vivo datasets. We chose simulation parameters for the synthetic examples, where -based on prior sensitivity analyses studies [7, 30, 67]-reliable estimation would be expected given our acquisition protocol, and assessed the relative performance of SBI vs MCMC.

We considered both single-shell (Eq. 3) and multi-shell cases (Eq. 4) with up to *N* = 3 fibre compartments. The number of model parameters inferred in the multi-shell case was 2 + 3 * *N*, where *N* is the number of fibre compartments, plus the noise variance. I.e., 6, 9 and 12 parameters for the 1-, 2-, and 3-fibres multi-shell model, respectively. For the single-shell models we had 1 parameter less (there is no *d*_*std*_).

#### Acquisition-specific vs Acquisition-agnostic training

As the above approach returns trained networks for a specific set of bvals and bvecs, we also explored using an acquisition-agnostic basis set representation of the data as input to the networks, instead of using directly the dMRI signal attenuation. The signal attenuations were expressed using Spherical Harmonics (SH) [68] (for single-shell) and the MAP-MRI basis (for multi-shell) [69]. Given the number of volumes in the data, we retained the even SH coefficients up to order *L* = 6 (28 coefficients), and for the MAP-MRI up to radial order *r* = 6 (45 coefficients) with Laplacian regularization (no positivity constraints nor anisotropic scaling). Both signal representations were obtained using Dipy implementations [70].

#### Probabilistic tractography

Using the trained architectures, we drew 50 samples from the posterior distribution of parameters given the observed data. For the same data and model, we drew 50 (thinned) posterior samples using MCMC as well. Each provided voxel-wise orientation distributions that mapped uncertainty for the estimated parameters. We subsequently used the estimated posterior orientation samples in probabilistic tractography. We ran the standardised FSL-XTRACT tractography protocols [71] using the estimates of each method. These protocols included a range of white matter tracts (commissural, projection, association and limbic). The SBI-derived and MCMC-derived tracts were then directly compared. As both of them reflect stochastic estimation processes, we further explored how the agreement in tract reconstruction between SBI and MCMC compares with scan-rescan reproducibility in MCMC-based tract reconstruction, across the six available repeats.

## 3. Results

### 3.1. Single-fibre model

We performed initial evaluations in the simple case of a single-fibre model (*N* = 1 in Eq. 3 and Eq. 4). Fig. 2 shows example comparisons for the posterior mean and uncertainty estimated by MCMC and SBI, in both single-shell (B) and multi-shell (C) models. A default SBI training paradigm was used in these first comparisons, with the same priors as in MCMC. Representative examples of the marginal posteriors for *f*_1_ in different areas of the brain are shown in Fig. 2A, with very good agreement between MCMC and SBI in the means of these posteriors, and some small variation in their coverage. Overall, as can be seen in the whole-brain maps, the agreement between these two very different inference approaches is extremely high, with correlations of the posterior means above 0.95 for both diffusivity and volume fraction, and median differences of 2^*o*^ for fibre orientations across white matter. Similarly, uncertainty maps show correlations of 0.92, i.e. a similar uncertainty mapping across the brain (lower uncertainty in WM and higher in CSF and GM for the fibre orientation). A very similar picture is depicted in Fig. 2C for the multi-shell model. These first findings suggest high agreement in both accuracy and precision of SBI estimates against MCMC, in the case of a simple model with a default training paradigm. In the next sections, we increase the model complexity and evaluate how different SBI training and designs affect performance.

**Figure 2:**
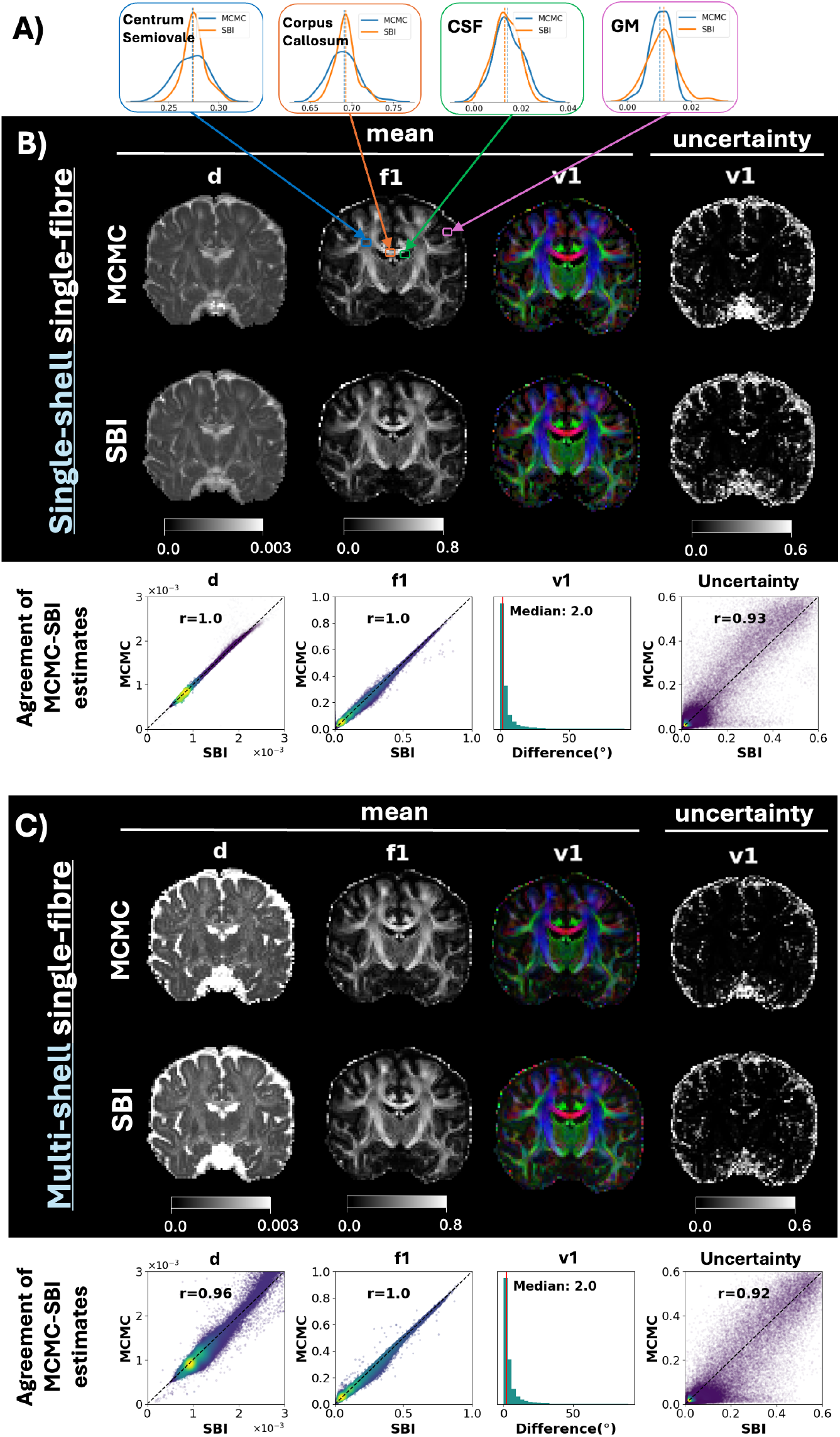
Comparison of SBI and MCMC estimates for a single-fibre model (*N* = 1) in case of single-shell. **(A,B) or multi-shell (C) data**. A) Example posterior density plots of the anisotropic volume fraction *f*_1_, as estimated by MCMC (blue) and SBI (orange), in four representative voxels: Centrum Semiovale (where fibre crossings are expected), highly anisotropic Corpus Callosum, CSF-filled ventricles, and gray matter. B) Whole-brain maps of estimated parameters using single-shell data (from left to right): Mean of *d* posterior, mean of *f*_1_ posterior, mean of ***v***_1_ posterior, orientation uncertainty (width of the ***v***_1_ posterior). C) Similar as in (B) for multi-shell data. For (B, C), density scatter plots represent the agreement of estimates throughout the brain; the correlation *r* is calculated for scalar parameters, and the angular difference (in degrees) is calculated for the mean fibre orientations. NPEs were trained with 1 million samples from uniform priors with random SNR values (within the range of 3 to 80).

### 3.2. Multi-fibre models - Synthetic data

#### 3.2.1. Optimal training strategies

We subsequently increased the complexity of the models to include crossing fibres (2 ≤ *N* ≤ 3). First, we compared different training strategies using synthetic data (where ground truth values for model parameters are known). We investigated 1) the increase in estimation accuracy due to usage of restricted priors compared to uniform priors, 2) the optimal strategy for considering noise during training, 3) whether alternative signal representations can provide similar performance (e.g., using the raw signal vs spherical harmonics), and 4) how different SBI designs perform (independent vs joint training). We generated data with *N* = 2 fibre compartments, where the anisotropic volume fraction *f*_1_ and the crossing angle were varied. Fig. 3 shows a collection of comparisons of the mean of the estimated (marginal) posterior distribution for volume fractions and crossing angles, demonstrating how accurately each inference approach captures the ground-truth patterns. For a representation of the error with respect to the ground-truth parameter values, see Suppl. Fig. S3; and Suppl. Fig. S4 for the precision/uncertainty in the estimates.

**Figure 3:**
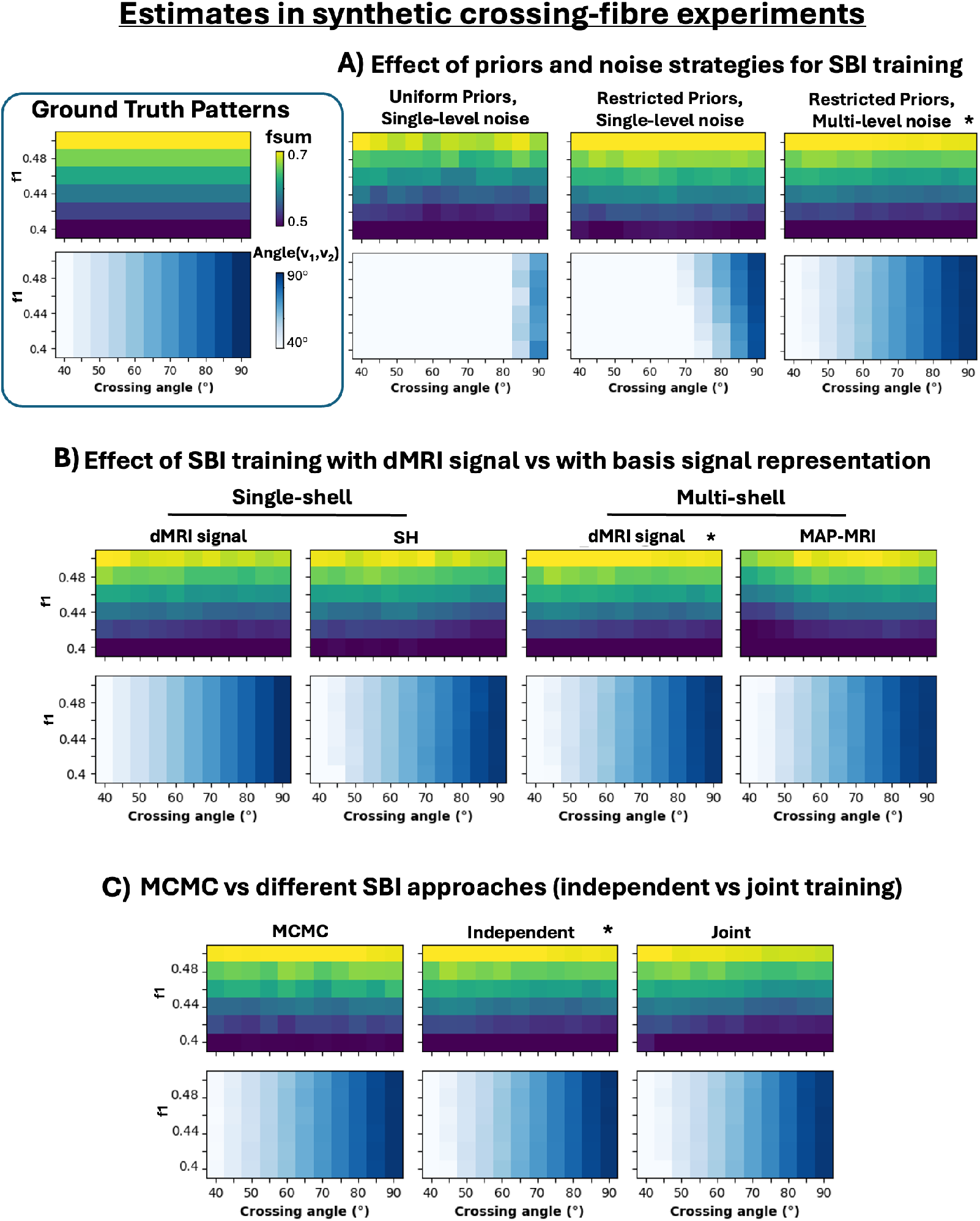
Evaluating performance of different S BI designs in estimating model parameters in synthetic fibre crossings (*N* = 2), where ground truth is known. dMRI was synthetically generated for patterns with two crossing fibres, varying the anisotropic volume fraction *f* _1_ and t heir crossing angle (rest o f parameters were *d* = 0.0012*mm*^2^*/s, f*_2_ = 0.2, *SNR* = 25 for all experiments). The ground-truth parameter values employed in these experiments are colour-coded (top-left corner), with each entry of these matrices corresponding to a different combination o f *f* _1_ and crossing angle. The ability to accurately estimate these values is explored with different S BI t raining designs. Estimates (median values over 100 noisy realisations) for the mean posterior of *f*_*sum*_ (top row) and crossing angle(*v*_1_, *v*_2_) (bottom row) are shown for the following cases: A) Different prior definition an d noise strategies for SBI training (from left to right): (i) Uniform priors and single-level noise, (ii) Restricted priors and single-level noise, (iii) Restricted priors and multi-level noise. B) SBI trained with raw dMRI signal (acquisition-specific) v s a basis set signal representation (acquisition-agnostic). Spherical harmonics were used for single-shell data, and the MAP-MRI projection for multi-shell data. C) Comparing MCMC fitting a model with *N* = 2 fibres against NPEs trained only with 2-fibres examples (Independent) or with a joint superset that contains both 1- and 2-fibres cases (Joint). All S BI models were trained with 5 million samples, apart from *Joint* approach which was trained with 6 million samples including 1- and 2-fibre cases (3 million each). Results denoted with (*) correspond to the same SBI design across A, B, C (i.e. trained with only *N* = 2 examples, multi-shell signal, restricted priors and multi-level noise).

Among the different strategies explored, using a multi-level noise scheme combined with restricted priors provided the best performance in retrieving the ground-truth. These improved both accuracy (see Fig. 3A) and precision (see Suppl. Fig. S4A). With restricted priors, the sampling efficiency was increased as all of the proposed parameter combinations met the conditions in Table 1. By sampling directly from “within-bounds” regions of the parameter space using the restricted prior, speed-ups of 62% were observed during the generation of the training sets, compared to sampling from uniform priors. This is because uniform sampling needed extra rejection steps to discard up to 39% of “out-of-bounds” parameter combinations (see Suppl. Fig. S2 for an example of how the parameter space gets constrained). Hence, multi-level noise and restricted-priors are used for any SBI model presented onwards.

Fig. 3B shows a comparison of the obtained estimates when SBI is trained against signal attenuation (for a given set of bvals and bvecs) or acquisition-agnostic signal representations, for both single-shell (using spherical harmonics) and multi-shell (using the MAP-MRI projections) scenarios. Some loss of accuracy can be observed in the signal representation cases, particularly at small crossing angles, suggesting that acquisition-agnostic training induces a small penalty in performance compared to training for data specific to a particular acquisition. However, overall good performance is preserved. For the rest of the paper we kept SBI trained against the full signal, but we will also present some in-vivo results using the MAP-MRI basis.

Finally, Fig. 3C shows the performance of MCMC (with *N* = 2 and no ARD priors) compared against NPEs trained only with 2-fibres examples (*Independent*) or with a joint superset that contains both 1- and 2-fibres cases (*Joint*). Both MCMC and *Independent* are always set to fit *N* = 2 fibre compartments (i.e. the right model for these data), while *Joint* inherently performs indirect model selection (it can fit either a *N* = 1 or *N* = 2 fibres model). Hence, the former corresponds to SBI_ClassiFiber in the ideal case of a perfect identification of the number of compartments by the classifier (N=2), while the latter corresponds to SBI_joint with *N* ≤ 2. Overall, all methods recovered appropriately the ground-truth values, with *Joint* having slightly worse performance than the other two, indicative of its higher complexity, but also its ability to indirectly perform model selection.

The above trends in terms of accuracy for the parameter estimates also hold in case of synthetic data where *N* = 3 fibres have been simulated (see Suppl. Fig. S5). Taken together, the findings suggest that SBI with appropriate training can perform very well in terms of posterior accuracy and precision (and similarly to MCMC) in synthetic data, when the model complexity (i.e. number of compartments) is a-priori known.

#### 3.2.2. Dealing with model selection

We subsequently explored more comprehensively the performance of different inference approaches with respect to model selection in 3-way crossing patterns. One of the main challenges when performing inference for a multi-compartment model is the identification of the number of compartments that are relevant to the observed signal. We explored two alternatives to deal with this challenge, SBI_ClassiFiber and SBI_joint. SBI_ClassiFiber performs an initial classification step to decide the appropriate model complexity *N* and then fits the respective NPE with *N* = 1, *N* = 2 or *N* = 3 compartments, each of which are independently trained. SBI_joint aims to learn the inherent structure by being trained jointly against *N* = 1, 2, 3 cases, avoiding the classification step and the need of training different NPEs. We compared these architectures in synthetic data against MCMC, for which we fit the most complex model (*N* = 3) and used ARD shrinkage priors for the volume fractions of each compartment [7].

Fig. 4A shows the performance for similar synthetic examples as before, where 2-fibre crossings were simulated, but up to 3 compartments could be estimated by all approaches. This allowed us to explore whether underfitting/overfitting occurred, contrary to the previous tests, where the number of compartments in the model was pre-fixed to the actual number of compartments simulated. All methods correctly identified the correct number of fibre compartments and showed good overall performance, with MCMC providing the best mean estimates. SBI architectures were close in estimation accuracy, with SBI_joint, as the most complex NPE, having slightly lower performance.

**Figure 4:**
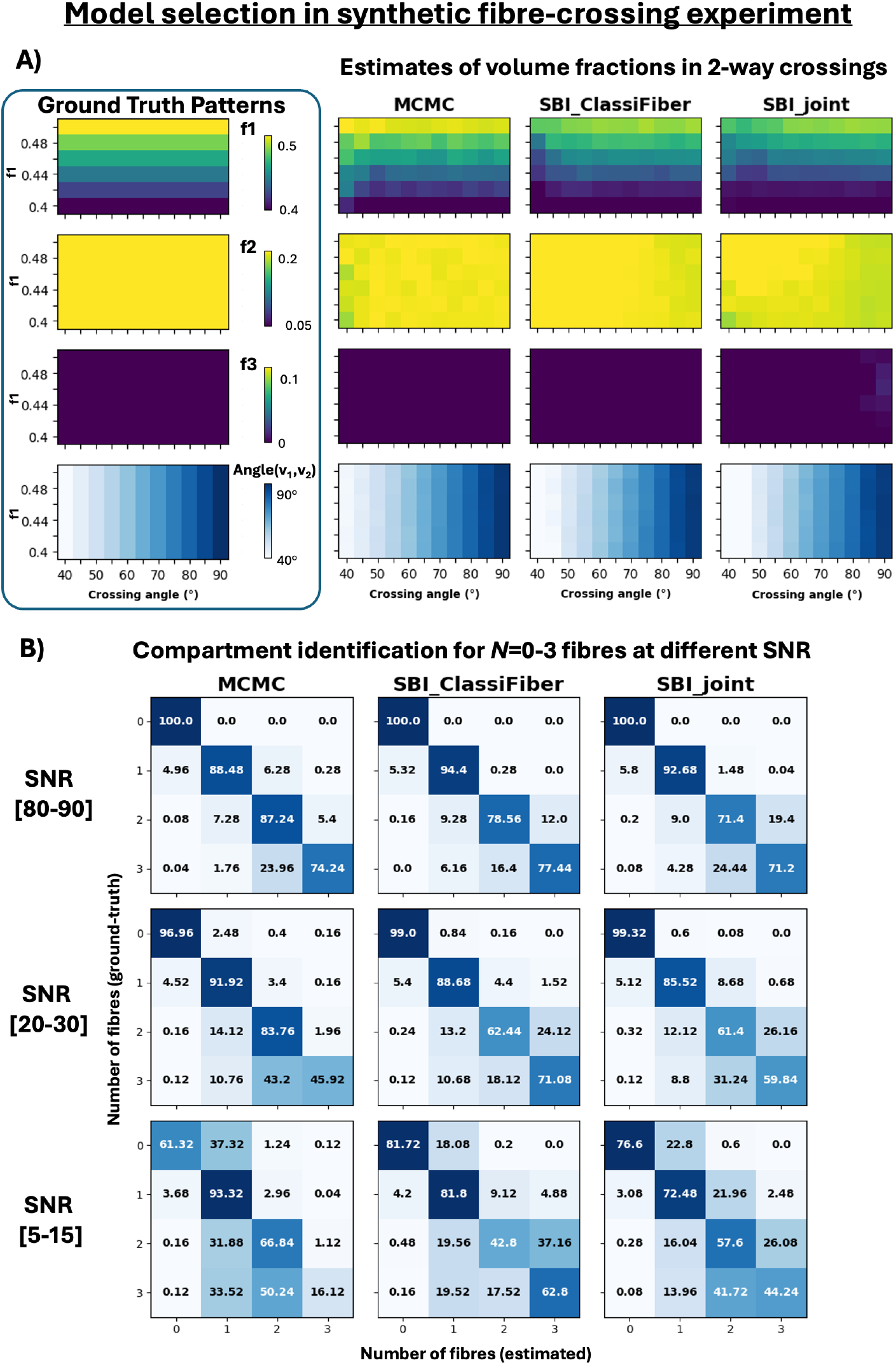
Evaluating performance in detecting the right number of fibre compartments in synthetic experiments, of MCMC, SBI_ClassiFiber and SBI_joint, all fitting up to *N* = 3 fibre compartments. A) Recovering a two-fibre pattern (same synthetic data as in Fig.3, where *f*_1_, *f*_2_ ≠ 0 and *f*_3_ = 0), when fitting 3-fibre models to detect over/under-fitting. Estimates (median values over 100 noisy realisations) for the mean posterior of *f*_1_, *f*_2_ and *f*_3_ and crossing angle(*v*_1_, *v*_2_) are shown. Both MCMC and SBI return zero *f*_3_ in all cases, detecting two fibre compartments with the correct volume fractions. B) Performance in compartment identification as a function of SNR for the three different approaches and for 10,000 synthetic patterns containing *N* =0-3 fibres compartments (with minimum crossing angle of 40^*o*^). For each SNR, matrices show for each ground-truth number of fibres (matrix row), the percentage of cases that were estimated as supporting *N* = 0 − 3 patterns (matrix column), i.e. each row of each matrix sums up to 100%. The ideal detection rate corresponds to a diagonal matrix; off-diagonal elements indicate over/under-fitting (above and below diagonal, respectively). In all cases, mean posterior *f*_*n*_ >5% was used to indicate support for the existence of a compartment. Results for SBI_ClassiFiber refers to the number of fibres supported by the final NPE estimates (i.e. *f*_*n*_ >5%), and not to the initial output of the classifier, which can be refined by the NPE.

We then tested the model selection performance in a larger set of 10,000 synthetic scenarios sampled from the restricted priors, with *N* = 0 − 3 (i.e. from zero to three fibre compartments, with various combinations of diffusivities, volume fractions and crossing angles). Fig. 4B shows in matrix format, comparisons of the number of fibre compartments detected (matrix rows) when different number of fibres were simulated (matrix columns). These detection matrices are presented for different noise levels and for the different inference approaches; diagonal matrices correspond to ideal model selection, while elements above/below the diagonal correspond to over/under-fitting. Overall, the three approaches achieved comparable results, with better detection rates (diagonal of the true vs predicted matrices) at higher SNR and simpler fibre configurations. At mid- and low-SNR, MCMC achieved a better identification of the 2-way crossings while SBI approaches provided a better detection of 3-fibre patterns, especially for SBI_ClassiFiber. At low-SNR, MCMC had the tendency to underfit (lower diagonal of the matrices) rather than overfit, which was not the case for the SBI approaches. This could be explained by the usage of a shrinkage ARD prior in MCMC [7], while the model selection in SBI is learnt directly from data.

### 3.3. Multi-fibre models - In-vivo data

After evaluating and optimising the performance of SBI in terms of accuracy, precision, and model selection in synthetic data, we compared them against MCMC using in-vivo dMRI multi-shell brain data. As there is no ground-truth available in the case of real data, MCMC estimates can be only treated as a reference. Fig. 5A shows a comparison of whole-brain mean and uncertainty maps obtained by the different methods for a model with *N* = 2 fibre compartments. Both SBI models provide very high correlations with MCMC (above *r* = 0.9 for all scalar parameters, see Fig. 5B). This inherently implies also a good agreement in the model selection results. Estimated crossing fibres show a good agreement as well (Fig. 5C and Suppl. Fig. 6A), with the median angular difference between SBI and MCMC being 2^*o*^ for **v**_1_ and 6^*o*^ for **v**_2_ in SBI_ClassiFiber (3^*o*^ and 8^*o*^, respectively, in SBI_joint). The example through the centrum semiovale shown in Fig. 5C, highlights all approaches resolving crossings between callosal and corona radiata projections, with the SBI architectures returning less of fibre fanning within the corpus callosum. Regarding orientation uncertainty, MCMC patterns (e.g. lower orientation uncertainty in WM) are overall reproduced through SBI although some broader posteriors were observed, especially in the second fibre orientation (see last two columns of Fig. 5A and B). Uncertainty maps for the secondary orientations have a bimodal distribution (low uncertainty for the voxels that support 2-way crossings and high for the rest in WM, GM and CSF). SBI approaches provided less “binary” uncertainty maps than the MCMC, with more regions having intermediate values between the two modes (i.e. between 0 and 0.6 - see Suppl. Materials for explanation of these values and Suppl. Fig. S7A for a different representation of these uncertainty plots).

**Figure 5:**
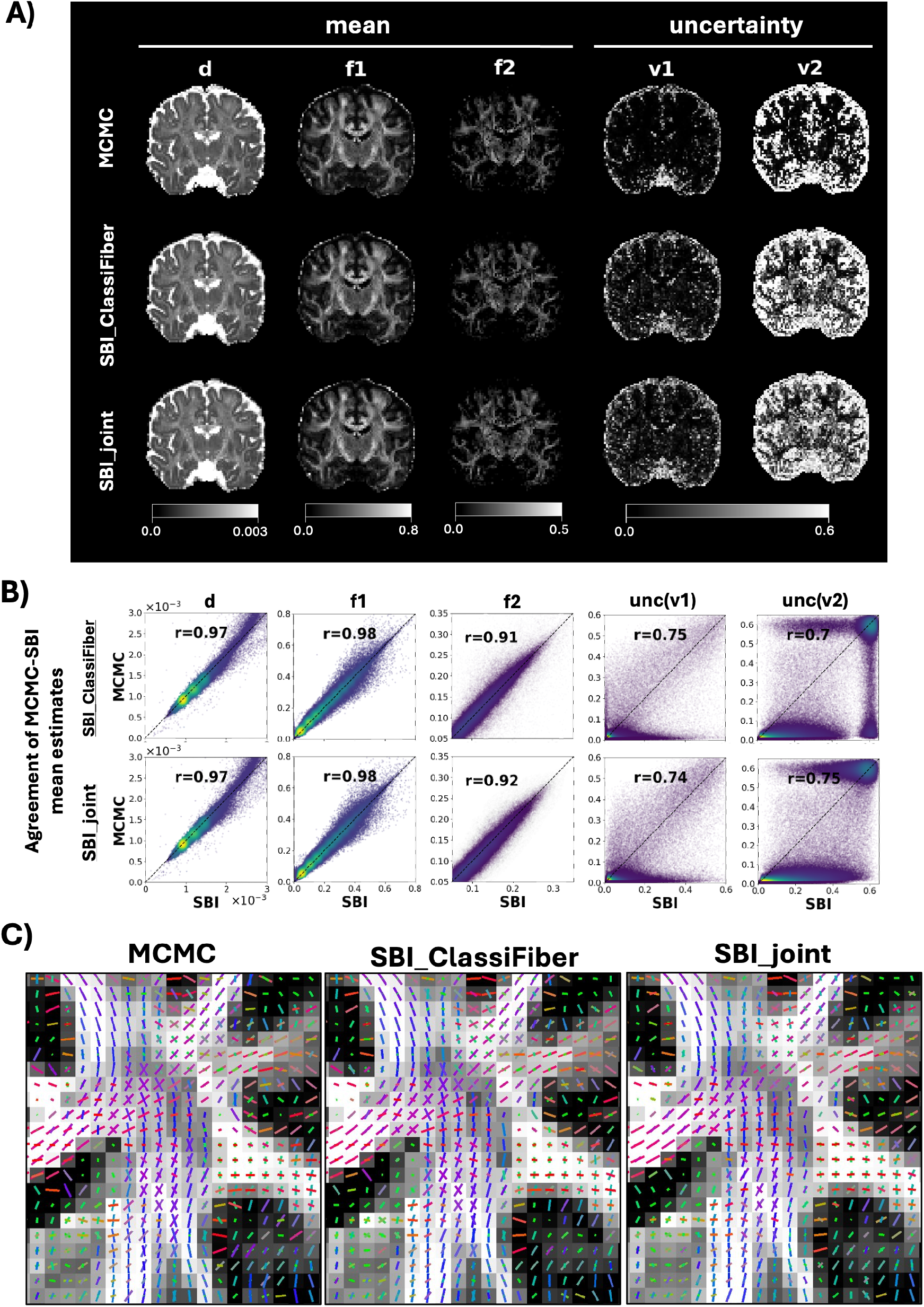
Inference for in-vivo brain data using a *N* = 2 fibres multi-shell model and different inference approaches (MCMC, SBI_ClassiFiber, SBI_joint). A) Exemplar coronal slice with the mean and uncertainty maps estimated by the different methods for different model parameters. B) Density plots representing the agreement between MCMC and SBI in the white matter; correlation *r* is calculated between scalar maps. C) RGB colour-coded crossing-fibres estimated by each method in the centrum semiovale. The mean posterior orientation is plotted for each compartment. The length of the orientation vectors are scaled by their relative voxel-wise volume fraction.

Similar trends can be observed in Fig. 6 for a *N* = 3 fibres multi-shell model. Panel A) shows mean maps of the diffusivity and volume fractions. As in Fig. 4, SBI_joint showed a more similar number of complex fibre patterns detected to MCMC, with 22.5% and 25.4% of the voxels in the white matter identified as 3-way crossings by each method, respectively, while SBI_ClassiFiber returned 33.3%. High agreement with MCMC was found for the first two compartments by both SBI methods, with correlations of *r*_*d*_ = 0.97, 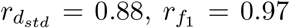, and 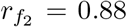 (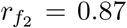 for SBI_joint). Larger differences were found in the third compartment; the correlation in the mean *f*_3_ reduced to 0.7 in SBI_ClassiFiber, and 0.58 in the SBI_joint.

**Figure 6:**
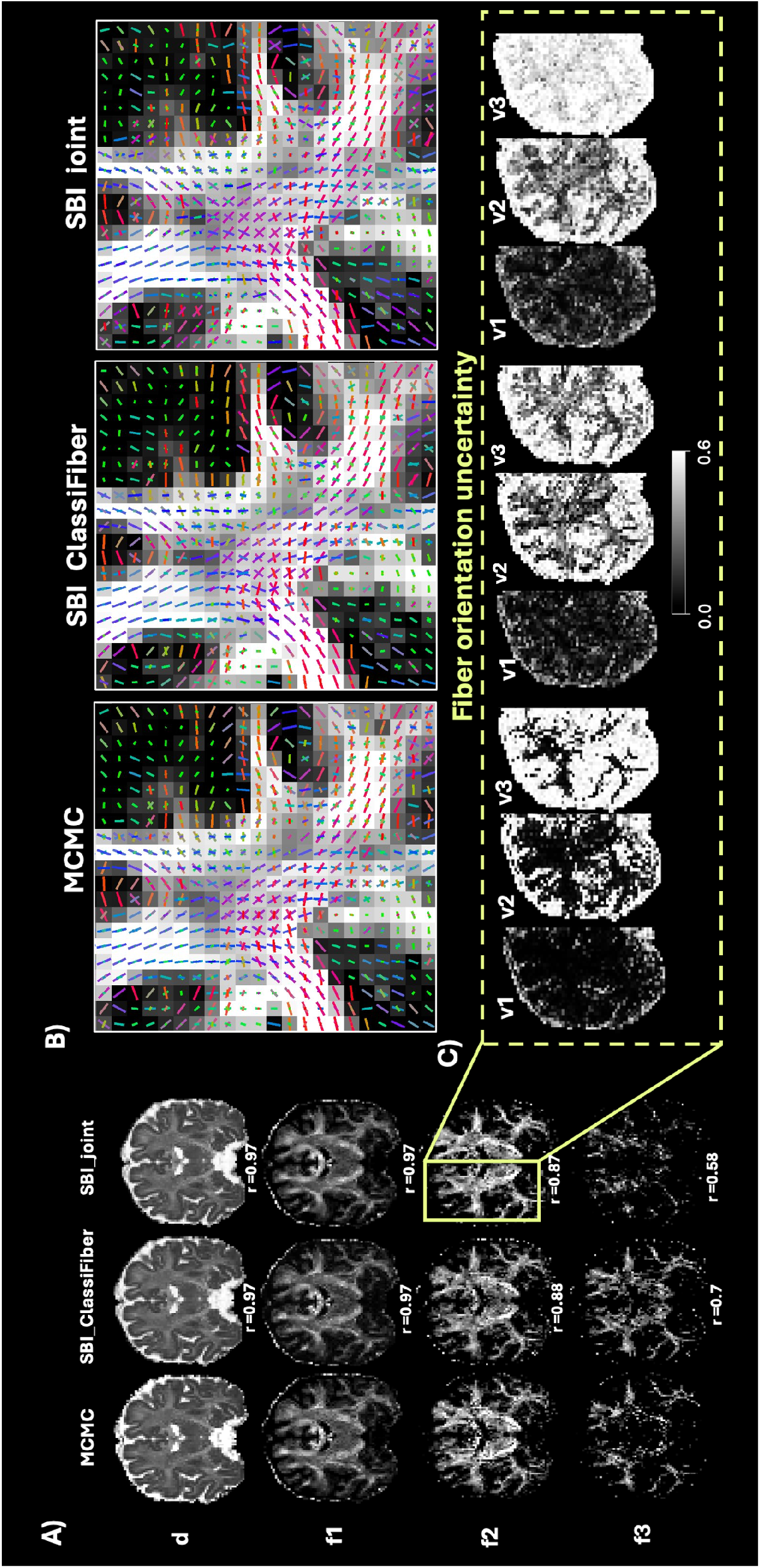
Inference for in-vivo brain data using a *N* = 3 fibres multi-shell model and different inference approaches (MCMC, SBI_ClassiFiber, and SBI_joint). A) Exemplar coronal slice with the mean scalar maps estimated by the different methods for different model parameters (correlation with MCMC is indicated at the bottom of each map). B) RGB colour-coded crossing-fibres estimated by each method in the centrum semiovale. The mean posterior orientation is plotted for each compartment. The length of the orientation vectors are scaled by their relative voxel-wise volume fraction. C) Orientation uncertainty maps for each fiber compartment and for each inference approach.

Panel B of Fig. 6 shows an example case of the orientations in the centrum semiovale, where 3-way crossings can be expected (corpus callosum fibres running left-to-right, corona radiata running superior-to-inferior and association fibres anterior-to-posterior). All methods depicted these, with SBI_joint having slightly lower coherence and crossings overall. Whole-brain orientation estimates are shown in Suppl. Figs. S6B.

Uncertainty maps for all three fibre orientations are shown in panel Fig. 6C, where SBI methods exhibited in general higher uncertainty compared to MCMC. Even if the general patterns were similar in all methods (i.e. highest uncertainty in CSF and GM, lower uncertainty in WM), SBI methods returned higher uncertainty in WM with increased model complexity.

Finally, as suggested in the previous section, using the MAP-MRI signal representations for a 3-fibres model led to similar parameter estimates (mean posterior) as using the raw dMRI signal (see Suppl. Fig. S8).

#### 3.3.1. Sources of SBI vs MCMC uncertainty quantification differences

So far we have observed that MCMC and SBI return similar patterns in the parameter estimates, but some discrepancies in the width of the posterior can be observed, particularly evident in in-vivo data. We further explored these differences in uncertainty quantification and identified scenarios where these occur.

A first identified case corresponds to the potential of fitting fewer compartments than supported by the data (e.g. due to inaccurate model selection). Fig. 7A shows the marginal and pairwise joint distribution for all parameters in an example synthetic case where a 3-way crossing pattern is simulated (crossing-angles of 90 degrees, *f*_1_ = 0.4, *f*_2_ = 0.3, *f*_3_ = 0.2), but a 2-fibre model is fitted, i.e. wrong model identification. MCMC returns unimodal marginal distributions for all parameters; it finds one of the 3 possible orientation modes and keeps sampling from it (as it is very *expensive* to jump to a different mode), missing the other modes and providing very low uncertainty (overconfidence). On the other hand, SBI captures all three orientation modes, assigning in this case their samples to (mainly) *v*_1_ (see red arrows) which, therefore leads to increased uncertainty due to multiple modes arised from fitting a model with the wrong number of compartments (epistemic uncertainty).

**Figure 7:**
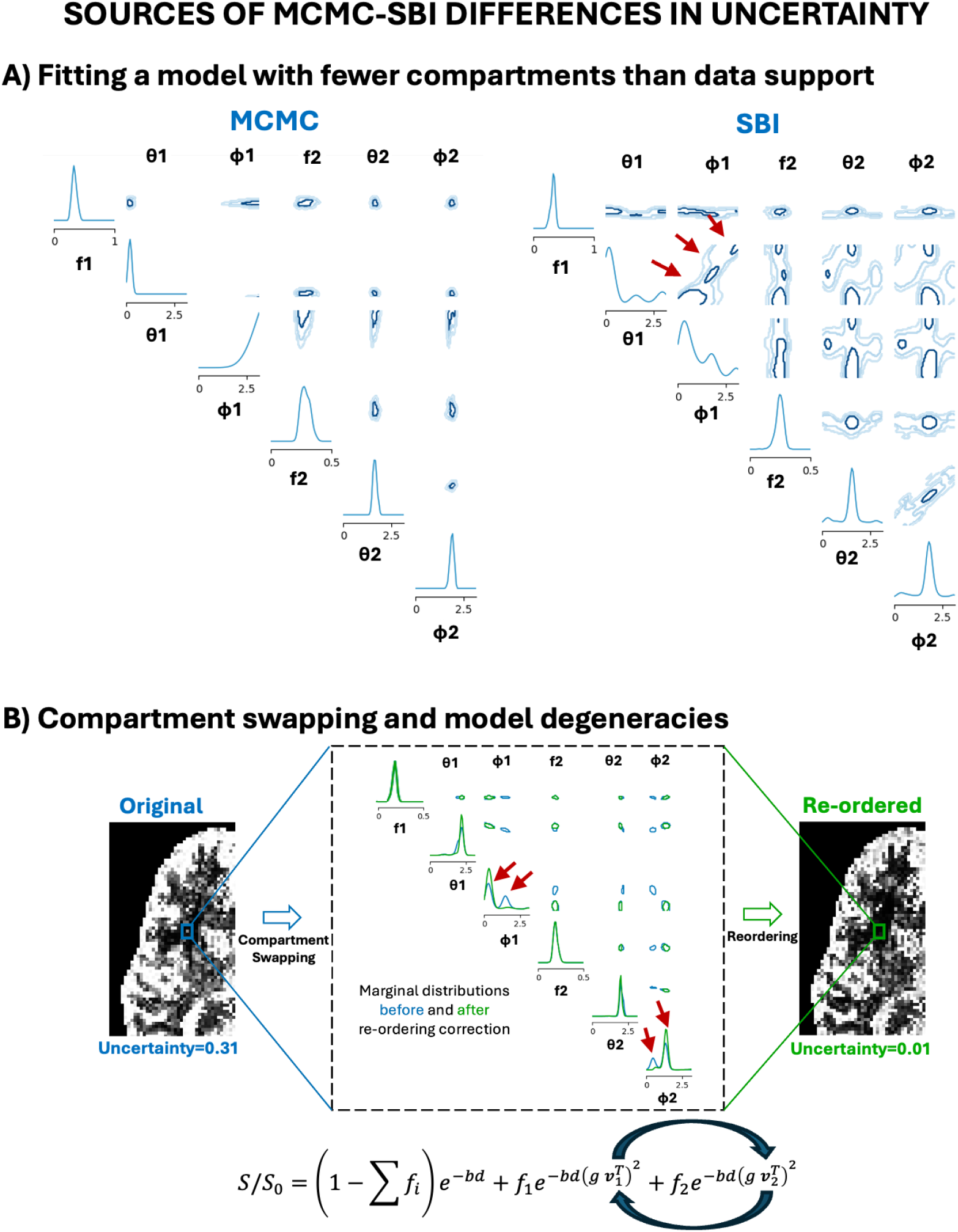
Sources of differences in uncertainty mapped with MCMC and SBI. A) Fitting fewer compartments than supported by data: Marginal and pairwise joint distributions of all parameters obtained from a 2-fibre model fitted to synthetic data from a 3-fibre pattern (ground-truth), using MCMC (left) and SBI (right). The marginal posterior distribution for every model parameter is shown first in each row, followed by the joint 2D posterior with every other parameter. When the number of compartments in the model is lower than the number of compartments supported by the data (i.e. effectively a wrong model is fitted), MCMC tends to return unimodal posteriors, missing the third compartment that is supported by the data. This leads to narrow (but wrong) distributions (overconfidence). SBI correctly captures all three modes within the compartments of the model (see red arrows, all captured in the **v**_1_ posterior in this example). However, this leads to broader posteriors (i.e. uncertainty). B) Compartment swapping: Marginal and pairwise joint distributions of all parameters obtained from SBI for a voxel from in-vivo data (square), where *f*_1_ ∼ *f*_2_ (blue). In such cases, SBI can include samples from the first and second fibre orientation indistinctively to the corresponding marginal posteriors, making them multimodal (red arrows). A simple re-ordering (based on similarity of orientation samples) can mitigate this (green), leading to unimodal posteriors. Notice that there is nothing wrong with either approaches, as the joint orientation posterior samples stay the same, and what changes is how they are distributed to compartments. For reference, orientation uncertainty maps of **v**_2_ returned by SBI, before (left) and after the correction (right) are shown.

Another source of differences in uncertainty is compartment swapping, which can occur when volume fractions of fibre compartments are very close to each other e.g. (*f*_*i*_ − *f*_*j*_) ≈ 0. In such cases, assigning a compartment to be first, second or third becomes arbitrary, as any combination could explain the signal in the same way. SBI is based on minimising the negative log-likelihood, so when the compartments have similar volume fractions, it will equally likely assign a sample from orientation **v**_*i*_ to parameter **v**_*j*_ and vice versa. Hence, drawing multiple samples can lead to a multi-modal marginal posterior for a specific fibre orientation parameter with *P* modes, with *P* the number of compartments with similar volume fraction. This compartment swapping is much more difficult to occur with a random-walk Metropolis Hastings MCMC algorithm, which needs very high energies to jump between multiple modes and therefore samples tend to inherently cluster together. Even if this issue in SBI is reduced by the prior constraint (*f*_*i*_ − *f*_*j*_ ≥ 0.05), it can still occur for relatively similar volume fractions that are just above this difference threshold. It can be further mitigated with a simple re-ordering of the posterior samples based on the similarity of orientations (rather than the default ordering based on the rankings of the volume fractions). Fig. 7B shows the marginal and pairwise joint distribution for all parameters in an example case from in-vivo brain data, with compartment swapping, before and after the re-ordering to compensate for the swapping. Suppl. Fig. S9 shows how reordering the SBI samples achieves very good correspondence between SBI and MCMC in an example voxel with 3-way crossings.

The above findings suggest that the differences observed in the uncertainty maps are potentially inflated by the way parameter samples are grouped into compartments and uncertainty is estimated. They also hint that individual samples in SBI are likely correct in all cases (Suppl. Fig. S9), even if summary uncertainty maps suggest otherwise when compared against summary maps from MCMC. We therefore performed probabilistic tractography, which uses the posterior orientation samples rather than summary uncertainty maps, to test this hypothesis.

### 3.4. Probabilistic tractography using SBI estimates

Probabilistic tractography spatially propagates orientation uncertainty (i.e. samples from the marginal posterior distribution of the fibre orientations) to build spatial distributions that estimate white matter pathways. We explored how the orientations estimated from the 3 different approaches (MCMC, SBI_ClassiFiber, SBI_joint) affect tractography performance. We used the posterior fibre orientation samples estimated by each method using a *N* = 3 fibres, multi-shell model. Fig. 8 shows example Maximum Intensity Projections (MIPs) of the path distributions for a range of 42 commissural, association, limbic and projection tracts [71]. For reference, the population-average UK-Biobank atlas is also shown, which was created using the same MCMC implementation for estimating fibre orientations and the same FSL-XTRACT implementation for tractography [71], as used here. All methods reconstructed all tracts successfully and showed a high qualititative agreement between the corresponding paths. SBI-reconstructed tracts had a higher correlation with the atlas (up to 15% more in SBI_ClassiFiber), compared to MCMC-based reconstructions.

**Figure 8:**
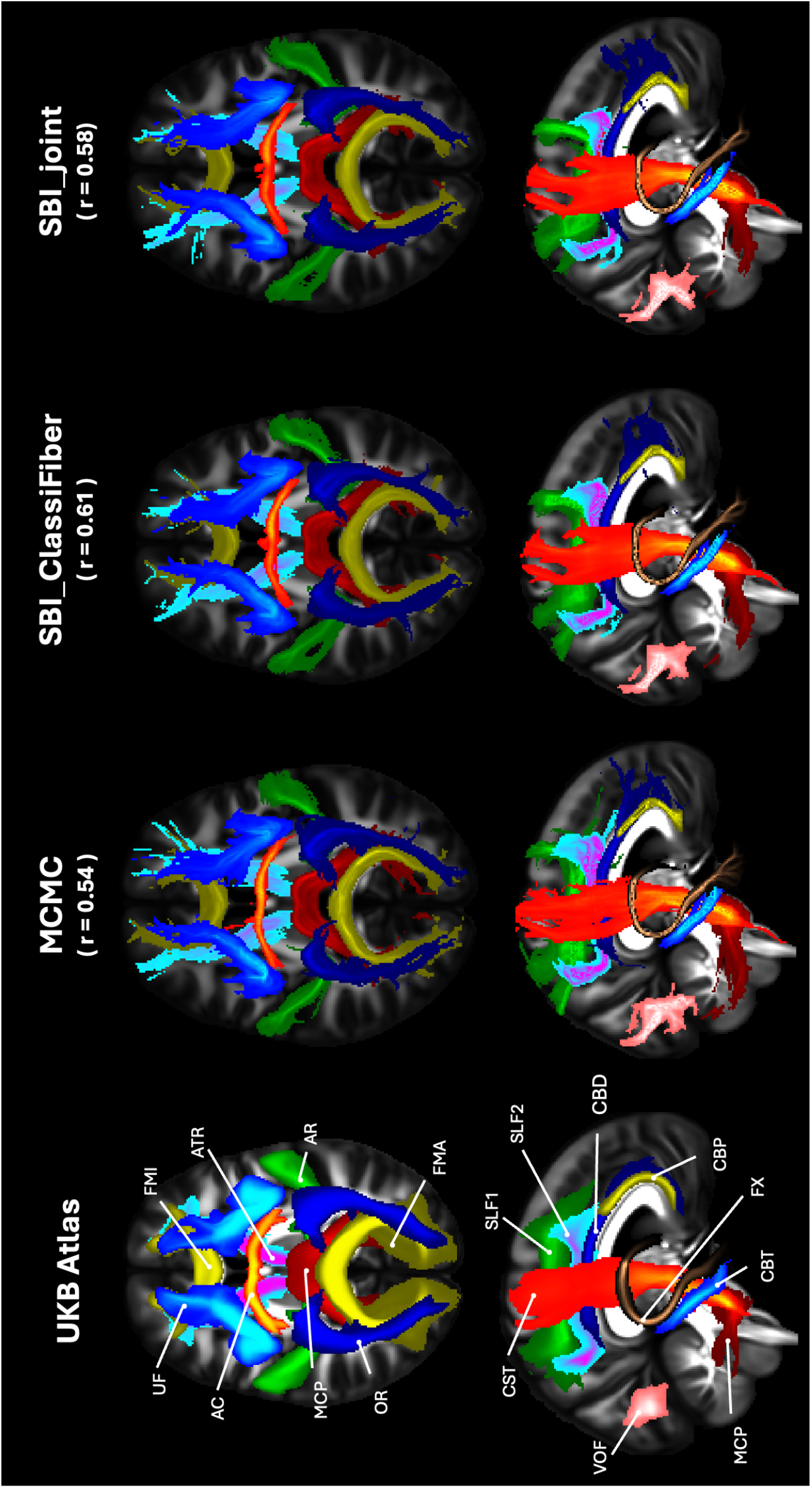
Probabilistic tractography using MCMC and SBI posterior orientation distributions. White matter tract reconstruction for 42 bilateral bundles were obtained using the orientation estimates of each method. The spatial distribution of each of the paths is shown as a maximum intensity projection on axial (top row) and saggital (bottom row) planes (thresholded at probabilities*>*0.1%). Tract reconstructions from a population-average atlas (using data from UK-Biobank and MCMC-based orientation inference) is shown as a reference. Each tract was spatially correlated against the corresponding reconstruction from the atlas. The median of these correlation values across tracts is indicated under each method.

We then looked into quantitative comparisons between the reconstructed tracts obtained using MCMC vs SBI estimates. Fig. 9A shows scan-rescan reproducibility of each method among 6 consecutive repeats of the same MRI acquisition protocol on the same participant. SBI methods reported higher reproducibility in tract reconstructions, with an overall median correlation *r* of *r* = 0.96 for SBI_joint and *r* = 0.95 for SBI_ClassiFiber (among tracts and repeats), which is consistently higher than in MCMC (*r* = 0.87). This provides evidence for the hypothesis that posterior orientation samples from SBI are appropriately precise and support the sources of the seemingly higher orientation uncertainty shown in the previous section. In addition, Fig. 9B demonstrates the median (among 6 repeats) correlation between MCMC-based and the corresponding set of SBI-based reconstructed tracts. This is overlayed on a shaded zone (in light blue), which depicts the interquantile range (across the 6 repeats) of the correlations between MCMC-based tract reconstructions themselves. In general, both SBI_ClassiFiber and SBI_joint perform similarly well to MCMC; certain tracts that have multiple 3-way crossings along their route are an exception (AF, AR, SLF1, SFL2, SLF3) as for these SBI_ClassiFiber performs better than SBI_joint. Nevertheless, these plots suggest that differences between MCMC and SBI in tract reconstructions are within the range of (and most of the time smaller than) scan-rescan variability (i.e., the variability of MCMC-based tract reconstructions across the 6 repeats). For the exceptions observed, where correlations between MCMC and SBI methods were lower, SBI methods showed a more precise reconstruction among repeats compared to MCMC (see 9A), as also indicated in Fig. 9C, where the averaged reconstructed tracts over the 6 repeats are shown.

**Figure 9:**
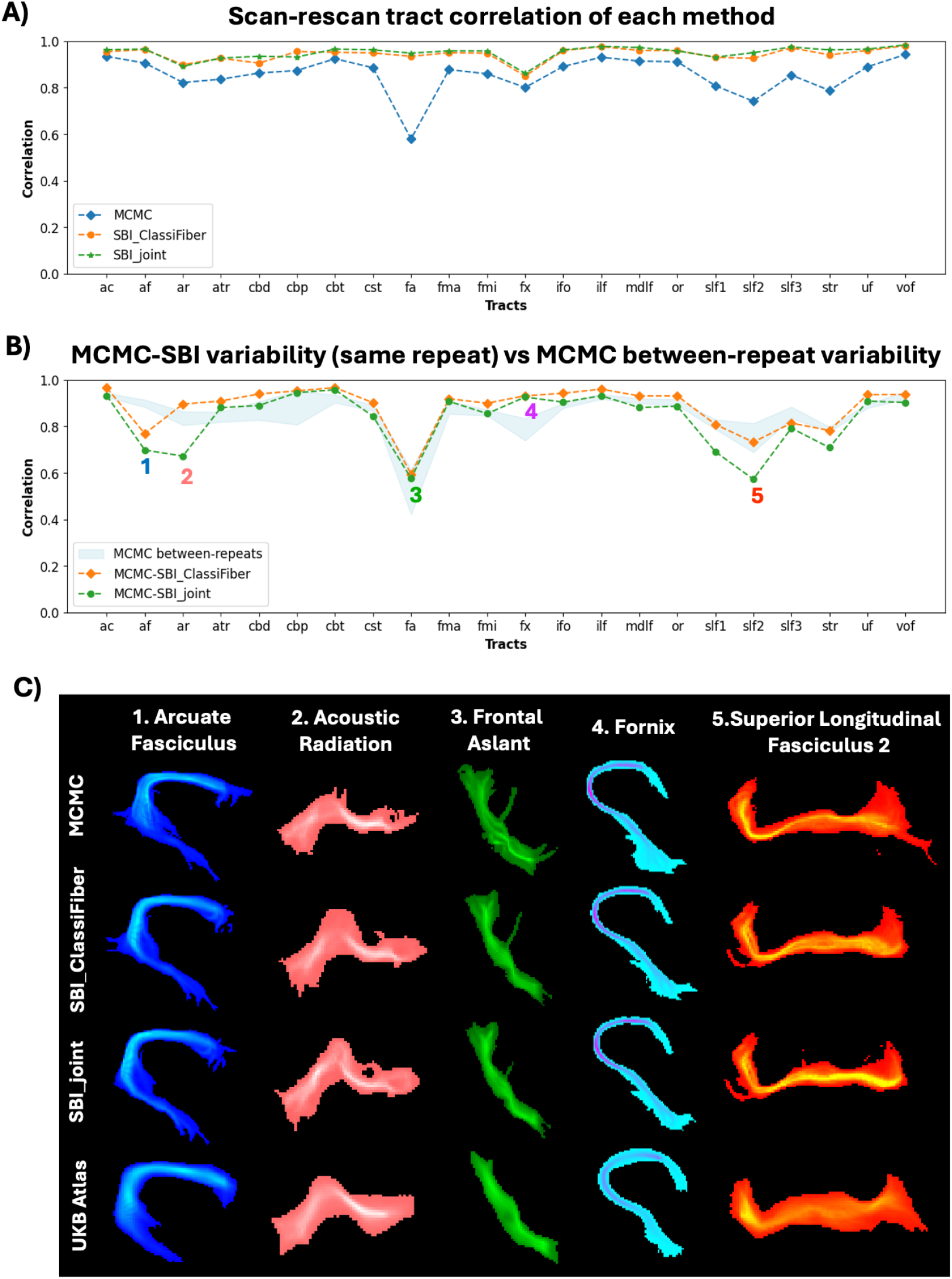
Scan-rescan reproducibility of probabilistic tractography reconstructions for a range of white matter tracts, across 6 MRI scan repeats of the same participant. Correlations of the spatial path distributions reconstructed for each tract and using orientation samples from MCMC, SBI_ClassiFiber and SBI_joint are shown. A) Median (across repeats) pairwise tract correlation for each method (i.e. median across rep1 vs rep2, rep1 vs rep3, rep1 vs rep4,… rep5 vs rep6). B) Correlation between SBI and MCMC-based reconstructed tracts (median across 6 repeats SB1 vs MCMC1, SB2 vs MCMC2,…) overlaid on the interquartile range of correlations of MCMC reconstructions across the 6 repeats. C) Averaged path distributions (across the 6 repeats) for a subselection of tracts that had lower agreement between MCMC and SBI.

Taken together, these results indicate that for a range of representative white matter tracts, including those with complex fiber architecture and 3-way crossings along them, SBI-based reconstructions i) agreed better with a population-average white matter atlas, ii) were more precise and reproducible across MRI repeats, iii) had differences with their MCMC counterparts that were within or lower than the magnitude of differences expected due to scan-rescan variability.

### 3.5. Computational performance

We measured single-CPU time required for simulation, training and parameter estimation in cases of *N* =1, 2, and 3 fibre compartments, and compared MCMC to SBI architectures. SBI has one-off overheads for the generation of the simulated training set and the training of the NPEs, but MCMC is considerably slower during inference. For the multi-shell models considered, the generation of synthetic signals with 3-way crossings took less than 30 minutes for 3 million samples. Time for SBI training increased with the number of fibres in the model and training samples, and both approaches required similar times. For instance, SBI_ClassiFiber with *N* up to 3 fibres required a total of ∼94 hours in a single CPU (∼2h for the ClassiFiber with 1M samples, ∼5h for the NPE with *N* = 1 and 2M samples, ∼33h for the NPE with *N* = 2 and 4M samples, and ∼54h for the NPE with *N* = 3 and 6M samples). Training SBI_joint with a single NPE but the same total number of 12M training samples with *N* up to 3 fibres required *∼*92 hours in a single CPU. However, the above can be significantly parallelised.

Inference times are compared in Table 2). Inference using the trained SBI networks was up to 70x faster than the MCMC, amortising the costs of training. We should point out that in these comparisons SBI is a CPU-based Python implementation, while the CPU-implementation of FSL-BedpostX used for these computational performance comparisons is highly optimised in C++. In summary, with a simple data-level parallelisation across e.g. 8 CPU-cores, inference on a standard in-vivo dMRI brain dataset, as the one used here, with the current SBI implementation takes ∼ 4 minutes, while MCMC takes more than 4 hours.

**Table 2:**
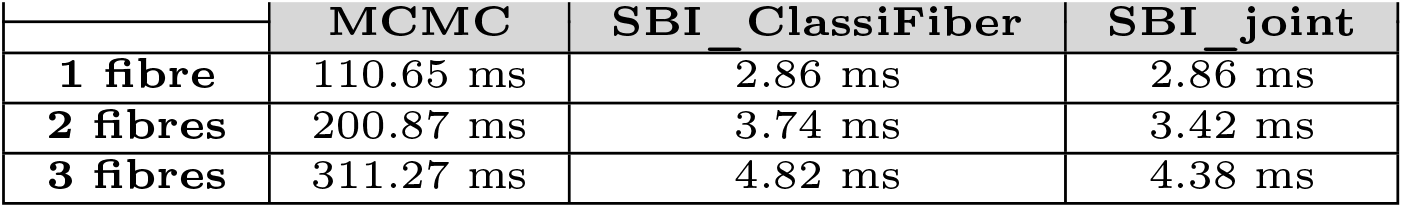
Computational performance - Inference time (in milliseconds) for each method for obtaining 50 samples from the posterior (reported times averaged across 1,000 synthetic cases). Times are all for a single CPU. Time for classification in SBI_ClassiFiber is orders of magnitude smaller and not included. MCMC includes 1,000 burn-in iterations and a thinning period of 25 (i.e. 2,250 iterations in total).

## 4. Discussion

In this work, we have presented a Simulation-Based Inference framework for Bayesian fitting of multi-compartment fibre orientation models in diffusion MRI data, mapping parameter uncertainty and utilising the posterior distributions of model parameters given the data to perform probabilistic tractography. Importantly, we performed a direct and comprehensive comparison against classical inference approaches, based on MCMC, to identify optimal training configurations and SBI architectures, and confirm the similarity of voxel-wise estimates and whole-brain tractography reconstructions to an established and heavily used approach. Our findings suggest that SBI with in-silico training can match the performance of classical inference approaches, opening new avenues for amortised inference in challenging likelihood-free scenarios [72]; while offering orders of magnitude of computational speed-up over MCMC and avoiding chain convergence challenges that MCMC can face.

We performed novel explorations for optimal SBI designs, in the context of dMRI modelling. Firstly, we explored how to incorporate noise into training. Previous studies have either not considered or added noise at random values [50, 52]. We have found that following a multi-level noise scheme improves accuracy and precision of SBI, even if 8-times fewer parameter combinations are considered for training (Fig. 3). From the perspective of neural networks applied to conditional density estimation, this noise-regularisation strategy can be seen as a form of data augmentation and is in line with recent findings [73].

Secondly, classical sampling-based, such as MCMC, can be very flexible in handling parameter constraints as prior knowledge. This is more challenging in an SBI framework, where parametric prior distributions are typically expected. We explored how to incorporate boundaries and non-linear constraints in the parameters. We showed that it is possible to train restricted prior objects to learn highly complex and non-linear constraints for the model parameters that translates into better performance (Fig. 3) and sampling efficiency (Suppl. Fig. S2), which can provide important speed-ups in problems where simulations are expensive or the parameter space is highly-dimensional. Density estimators can struggle to generalize across the entire parameter space even for simple problems [74], so a trade-off between enough diversity to capture the high-dimensional parameter space (restricted priors) and having enough, slightly-perturbed, samples (noise strategy) can be beneficial during training, as shown by our results.

Thirdly, we explored two strategies for handling model selection in cases of nested multi-compartment models. Recent studies have proposed a classification step that chooses what model complexity to fit [52, 54] followed by an NPE trained independently on that model. We took a step further, inspired by recent studies [75, 64], and evaluated our implementation of this approach (SBI_ClassiFiber) vs another one where the NPE learns the appropriate model complexity by being trained on multiple potential models (SBI_joint). Results in both synthetic and in-vivo brain data suggest a comparable performance of both approaches, with SBI_joint providing slightly worse crossing fibre rate detection, accuracy and precision than SBI_ClassiFiber, but tractography end results suggesting similar or better performance than MCMC (which relied on ARD shrinkage priors for indirect model selection [76]). We showed that training an NPE with enough examples from different models can be sufficient to learn the model selection, at the cost of some precision and accuracy loss. This is important, as in SBI_ClassiFiber (as well as in similar alternatives proposed, e.g. [64, 4]), the classifier and NPEs are trained independently, so a misclassification by the former will directly affect the latter without any feedback. However, by learning the complexity indirectly as part of training, model selection and parameter fitting can interact during inference. Exploring the joint-training of the ClassiFiber and the different NPEs to allow for such interactions could shed more light on what are the advantages and disadvantages of each architecture.

We evaluated the inference performance of SBI against an MCMC-based framework for the same models, which we have developed in the past [7, 31, 30, 24], and is widely and heavily used as part of FSL [77]. For both synthetic and in-vivo brain data, we found that SBI closely reproduces the patterns of MCMC. In terms of agreement in the means of the posterior distributions (as a proxy for accuracy), correlations above 0.9 between SBI and MCMC architectures were found across the brain for most of the parameters in the different models. In terms of uncertainty mapping, correlations above 0.9 were found for simple models (with a single fibre compartment, Fig. 2). As the number of compartments increased, the whole-brain patterns remained (i.e. maximum orientation uncertainty in GM and CSF, lower in WM) but orientation uncertainty estimated by SBI in the WM was generally higher than by MCMC (see Figs. 6 and Suppl. Fig. S7). Longer training did not change these trends considerably (results not shown). Despite these seemingly large differences in uncertainty mapping, probabilistic tractography (that spatially propagates uncertainty in orientation estimates) using MCMC and SBI approaches returned very similar results. In fact, SBI-based tract reconstructions agreed better with a population-level white matter atlas, were more reproducible across repeats of the same MRI experiment, and their differences with MCMC-based reconstructions had a magnitude smaller than expected from scan-rescan variability (Figs. 8, 9).

We explored what could drive this discrepancy between uncertainty mapping at the voxel vs whole-brain level. One clear difference is how uncertainty is depicted at the two levels: a) At the voxel-level, we used the approach implemented in FSL [7, 31] both for MCMC and SBI. This discretizes the joint posterior orientation distribution by grouping orientation samples into a finite set of *N* fibre compartments, and subsequently calculates their mean orientation and spread around it using vector dyadic products [5]. This provides a mean orientation and uncertainty around it for each fibre compartment supported per voxel. b) At the whole-brain level probabilistic tractography propagates using directly the orientation samples, i.e. from the joint posterior of orientations. Hence, if the grouping of orientation samples into finite sets is not consistent between MCMC and SBI, large differences can be observed at the voxel level and not at the tractography level. In Fig. 7 and Suppl. Fig. S9, we presented scenarios and confirmed with examples from the data where this occurred.

Fitting the wrong model to data can be one such scenario (see Fig. 7A). This adds epistemic uncertainty and increases the challenge of what the mapped uncertainty represents. Even if SBI and MCMC account explicitly for aleatoric uncertainty by considering the effects of noise in the training dataset (see Suppl. Fig. S10), epistemic uncertainty will be also indirectly captured in the estimated posteriors. However, this can be reflected differently in SBI and MCMC. Specifically, in cases of multi-modal orientation distributions, we observed how SBI could capture all the modes (even with a model with fewer compartments than modes), spreading the samples across modes and adding into orientation uncertainty. While the MCMC implementation sampled only from certain modes, providing samples with high precision, but completely ignoring an existing mode of the target distribution (Fig. 7A). Multiple modes can also arise from non-injective models, where different combinations of parameter values produce similarly likely solutions (i.e., degeneracies). This is a known challenge for certain microstructural models and the multi-modal posterior can be used to evaluate the different likely solutions, as illustrated in Jallais et al [50]. Here, by similar mechanisms, we have identified other type of degeneracies using SBI due to compartment swapping (Section3.3.1).

### 4.1. Limitations and Future work

Even if we aimed to match as closely as possible MCMC to SBI implementations in order to compare their performance, a perfect matching is not feasible as they represent quite different approaches. A number of algorithmic features can be contributing to observed differences. Firstly, the MCMC implementation relies on ARD shrinkage priors, which introduce large a-priori penalties on model complexity and have considerable effects on fibre crossing detection and on uncertainty mapping. ARD priors lead to more “binary” values on the uncertainty in the fibre orientations (either maximum or null uncertainty, see uncertainty maps in Figs.5 and corresponding histogram of Suppl. Fig. S7C). Such improper priors cannot be implemented for SBI in a straightforward manner. Secondly, SBI networks use a Kullback-Leibler divergence loss function to train NPEs. Due to the mode-covering property of the KL divergence, the approximate posterior will always cover the true posterior [78] with deviations leading to broader posteriors. In density estimators, this is known as *leakage* and is particularly problematic in neural spline flows because they do not have analytical corrections [38, 39, 79, 41]. Longer training or refinements, such as importance sampling have been proposed to compensate to some extent to converge to the true posterior [80, 81] but it is still an open problem that requires more research. Thirdly, the Metropolis-Hastings MCMC implementation used for comparisons is based on local sampling techniques. Hence, the energy needed to jump from one mode of a posterior to another can be very high. In this way, MCMC is more likely to converge to one of the modes and “stick to it” than SBI, missing other likely modes. This also suggests that SBI can perform a better global search than MCMC methods, which highly depends on the initialisation and the progression of the chain. Despite all these factors, the level of similarity between MCMC and SBI in local and global estimates was remarkable and considerably higher than the differences.

Another challenge is disentangling aleatoric from epistemic uncertainty in Bayesian approaches, and appropriate metrics are also needed for this purpose. We explored the ability of SBI to estimate the noise level in the data (i.e. noise variance), the main source of aleatoric uncertainty. Suppl. Fig. 10 demonstrates that estimation of noise levels is reasonable both in synthetic data and real data. It shows, however, how both MCMC and SBI can deviate from estimates that FSL-eddyqc [82] provides, which are obtained from the repeated *b* = 0 measurements directly in a model-independent way. As SNR estimates can become dependent on the model and noise model used (epistemic uncertainty), it is debatable on whether such estimates should be preferred upon measurements that can be directly obtained from the data. This is an interesting topic and future work is needed to identify what would be the optimal way forward.

In terms of performance, scalability with parameter space dimensionality can be an issue for all inference approaches. In this work, we explored multi-compartment models with different number of compartments and SBI performed as well as MCMC for tractography with the highest number of compartments typically considered. Recent studies have considered models with very high dimensionality and demonstrated that SBI can provide accurate solutions even for large parameter spaces [83]. However, orthogonality in parameter space may interact with scalability and further work is needed to explore these aspects comprehensively.

A limitation specific to ML-based frameworks such as SBI is that training against the dMRI signal involves model predictions for a particular acquisition scheme, i.e. a particular set of *b* values and diffusion-sensitising gradient orientations in the case of dMRI. Even if re-training for new data and acquisition schemes can be done once and is relatively cheap, another approach would be an model-agnostic representation of the the dMRI signal, e.g. by a number of orthonormal sets of basis functions or any other latent representation learnt by the networks [50, 52, 54]. This would allow acquisition-agnostic inference, as NPEs are trained against generalisable basis sets rather than signal sampled with a specific scheme. We explored training against such representations, including spherical harmonics (for single-shell) and MAP-MRI (for multi-shell data). Promising results in both synthetic and in-vivo brain data suggested that NPEs trained against both representations could provide similar accuracy performance compared to NPEs trained against the signal, at the cost of some precision loss (Fig. 3B and Suppl. Fig. S4). This would be interesting to explore further, where spatial neighbourhood information could be also explored for this purpose [84].

In summary, we argue that SBI is a powerful approach for performing Bayesian inference and has shown comparable performance to an established and heavily-used MCMC-based framework. Although further research and validation is needed, SBI could pave the way for tackling unaddressed problems in neuroimaging, given its reliance on in-silico, likelihood-free forward predictions and its scalability to multi-dimensional problems, the two main limitations of classical Bayesian inference approaches. The advances presented here on fibre orientation mapping are directly applicable to usage scenarios of tractography towards clinical translation, for instance in the development of structural scaffolds for whole-brain personalised models of neuronal dynamics in health [85] and disease [86] or in neurosurgical planning [87].

## Acknowledgements

JP and SS are supported by an ERC Consolidator Grant (101000969), and JP is also supported by a Wellcome Trust bioimaging technology award (313367/Z/24/Z). MD, CS and JHM by the German Research Foundation (DFG) through Germany’s Excellence Strategy (EXC-Number 2064/1, PN 390727645) and SFB1233 (PN 276693517), the German Federal Ministry of Education and Research (Tübingen AI Center, FKZ: 01IS18039) the Carl Zeiss Foundation and the Else Kröner Fresenius Stiftung (Project ClinbrAIn).

## Supplementary Material

### MCMC implementation

We used MCMC implementation of FSL-bedpostx [7, 30, 31]. The likelihood is using the Ball&Sticks forward model and is defined assuming: a) independence between diffusion-sensitizing measurements *j* = 1 : *J*, b) additive zero-mean Gaussian noise with precision *τ* = 1*/σ*^2^. Gaussian noise is a good approximation for Rician MRI noise when the SNR is larger than 3 [88]. The likelihood function, s*p*(*Y* |**Ω, *τ***) is therefore:

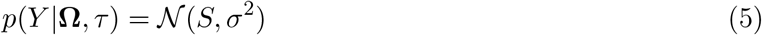

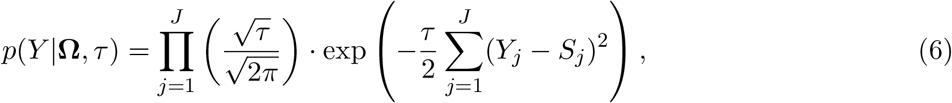

where *Y*_*j*_ is the measured signal with gradient *j*, Ω are model parameters of interest, and *S*_*j*_ is the predicted signal by the forward model, given by the Ball&Sticks model (see eq. 3 and eq. 4 in the main text). Noise variance is marginalised out using reference priors (so that it is not inferred), leading to a likelihood dependent only on model parameters Ω [21]:

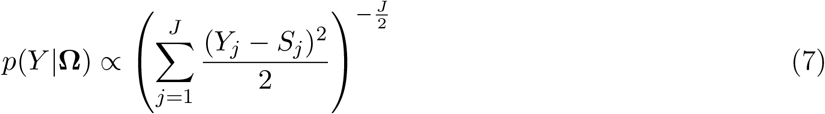

Regarding the prior distributions of the parameters, the prior on spherical angles ensures a uniform prior on the sphere, while positivity constraints are used on *S*_0_, *d*, and *f*_*n*_ [21].

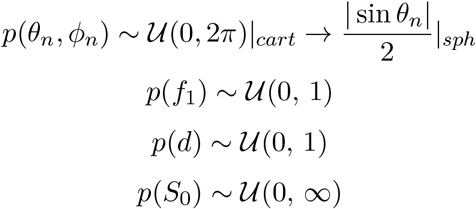

In case of multiple fibre compartments, the sum of all anisotropic volume fractions is constrained to be in [0,1]. In addition, Automatic Relevance Determination (ARD) priors are used for the corresponding volume fractions *f*_*n*_, *n* ≥ 2. ARDs are shrinkage priors that a-priori heavily penalise model complexity and the existence of extra compartments, providing high support for near to zero values for volume fractions *f*_*n*_, *n* ≥ 2. They therefore allow indirect model selection to be performed, when the most complex model is fitted, with low prior support for multiple compartments. As indicated in [7], ARD priors are *p*(*f*_*n*_) ∼ 1*/f*_*n*_ (*n* ≥ 2).

A univariate Random-walk adaptive Metropolis-Hastings algorithm is used for MCMC. Model parameters are initialised by a non-linear deterministic fit (using Levenberg-Marquadt non-linear optimisation). Proposals for MCMC are univariate Gaussians, centred in the last accepted sample and with adaptive variance to ensure a balanced acceptance/rejection ratio of 50%. MCMC runs for 1,000 burn-in iterations that allow the chain to converge and subsequently 1,250 samples are obtained for each parameter. To avoid autocorrelation in the Markov chain, one of every 25 samples is retained (thinning=25). Hence, after thinning and burn-in, the approximated posterior distribution of each model parameter contains 50 samples.

### Uniform sampling on the sphere

Linear and spherical geometries obey different principles. A uniform sampling in the plane does not return a uniform sampling in the sphere, as there is a different surface area for the same number of points. Close to the poles of the sphere, the differential surface area element gets smaller (it shrinks), and the wrapping of the uniform planar distribution produces an oversampled area. In contrast, close to the equator, the differential surface area increases (it stretches) and the density of points is lower (see Suppl. Fig. S1A). Thus, we should include fewer points near the poles and more points near the equator to achieve a uniform distribution on the sphere. We apply the following correction to achieve uniform sampling in the sphere when training and evaluating the SBI framework (see Suppl. Fig.S1B):

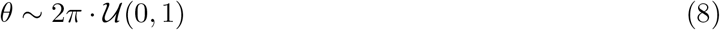

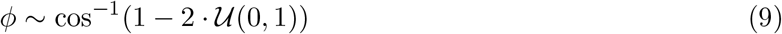

### Estimating mean fibre orientations and their uncertainty

In order to obtain the mean fibre orientation of *M* vector samples described by spherical angles (*θ, ϕ*), we use the average dyadic tensor [5]. After a spherical to cartesian coordinates conversion, the dyadic tensor corresponding to a vector 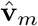 is the outer product of the vector with itself. The average dyadic tensor **V** is the mean over the M samples:

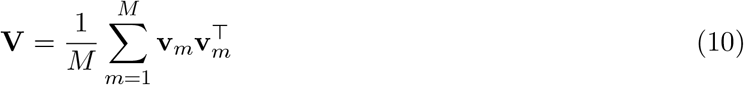

The principal eigenvector **e**_1_ of the mean dyadic tensor **V** (i.e. the one corresponding to the largest eigenvalue) represents the mean orientation of the set of *M* vector samples. The corresponding eigenvalue provides a measure of how concentrated the orientations are around that direction **e**_1_. Hence, the orientation uncertainty can be computed as:

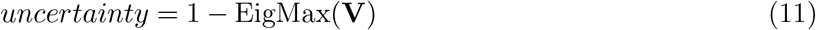

**Figure S 1:**
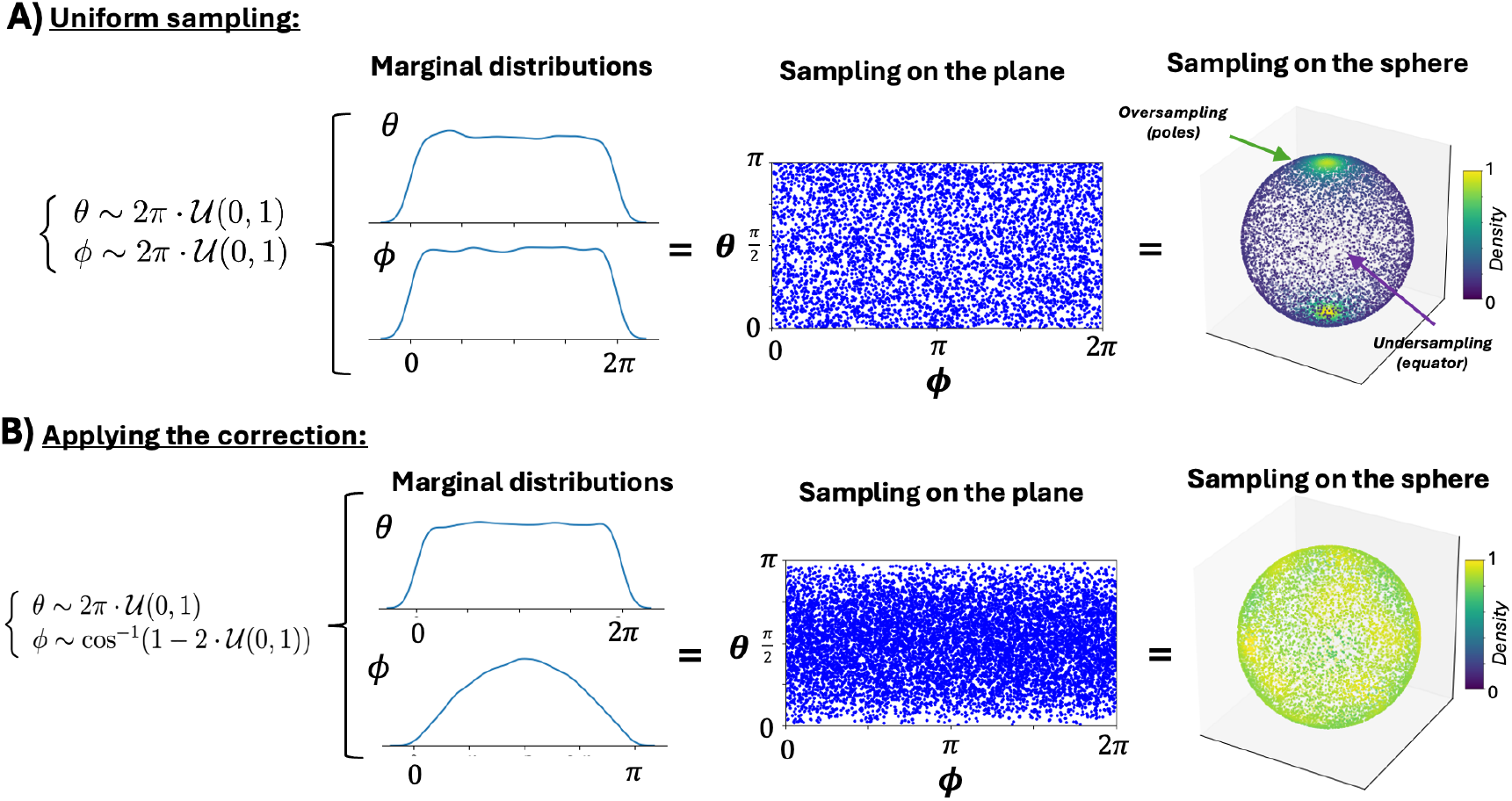
Defining uniform priors on the sphere for fibre orientations. A) Uniform sampling of orientations in the interval (0, 2*π*) leads to uniform coverage on the plane, but not on the sphere, as it produces oversampling at the poles and undersampling at the equator. B) Applying the correction, uniform sampling on the sphere can be achieved, even if this does not look uniform on the plane *θ* − *ϕ*. In both A) and B), from left to right: (i) Definition of the prior distributions of the fibre orientations, (ii) Marginal distributions produced when sampling from such priors, (iii) joint distribution represented on the *θ* − *ϕ* plane, (iv) joint distribution represented on the sphere, colour-coded by the density of the samples.

Since **v**_*m*_ are unit vectors, **V** is symmetric and positive semi-definite. The eigenvalues of **V** represent the spread or concentration of the orientation samples. If the orientations are perfectly aligned, EigMax(**V**) = 1, and the other eigenvalues are zero. This corresponds to no uncertainty (uncertainty = 0). On the other hand, for completely isotropic distributions (uniformly sampled over the sphere), the eigenvalues of **V** are equal, and the largest eigenvalue is 1/3. This gives the highest uncertainty (*uncertainty* = 1 − EigMax(**V**) = 1 − 1*/*3 ≈ 0.66).

### Accuracy and Uncertainty Metrics

To provide quantitative performance metrics, a reference value is generally needed. For comparison of the mean estimates in synthetic data, these reference values are the ground-truth parameter values used to simulated the dMRI signals. However, there is no ground-truth for uncertainty, nor for estimates in in-vivo brain data. In those cases, we compared the SBI estimates against the estimates from the established MCMC implementation in FSL-bedpostX, i.e. the latter are used as the reference values.

#### Accuracy

The agreement between scalar parameters is generally indicated as the linear Pearson’s correlation *r* between the reference values and the mean of the estimated posterior distribution samples, while errors are calculated as the % of their difference:

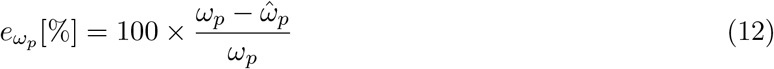

For fibre orientations, the angular difference ∠*e* was defined as the crossing-angle between the reference value **v**_*n*_ and the mean estimated 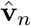 from the posterior samples:

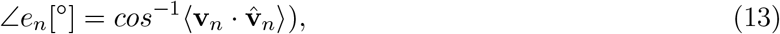

where 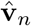 is the dyadic vector, i.e. the principle eigenvector of the dyadic tensor (see section above). In the case of multiple fibre orientations, we matched the closest pairs before calculating the errors per orientation.

For model selection, we calculated the confusion Matrix (CM) of the actual number of fibres *N* vs predicted number of fibres 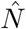 for each approach. This offered not only the detection rates for each fibre but also potential trends and patterns in the mis-classified cases.

For tractography evaluations, for each estimated tract, the corresponding spatial distribution was thresholded (≥ 0.001) to remove noisy tails. To compare a pair of corresponding tract reconstructions, we calculated the Pearson’s correlation between the respective spatial distributions (after vectorising each of them). To avoid bias from any of the tract reconstructions in comparison, the correlation was performed within a population-based atlas-derived tract mask, available from [71].

### Uncertainty

For scalar parameters, the standard deviation (width) of the posterior distribution was used to obtain a metric of uncertainty. For fibre orientations, the maximum eigenvalue *λ*_*n*_ of the dyadic tensor of the posterior orientation samples was used to provide a measure of uncertainty, i.e. how concentrated the orientations were around the mean direction of the samples. In synthetic data, we evaluated how such orientation uncertainty changed as a function of noise level and training data size, as uncertainty is expected to reduce as the SNR and/or the training data size increase (see also [55]). In real data, we used MCMC uncertainty maps as reference for direct comparisons, and the probabilistic tractography results as indirect indications of how consistent spatially-propagated uncertainty were.

**Figure S 2:**
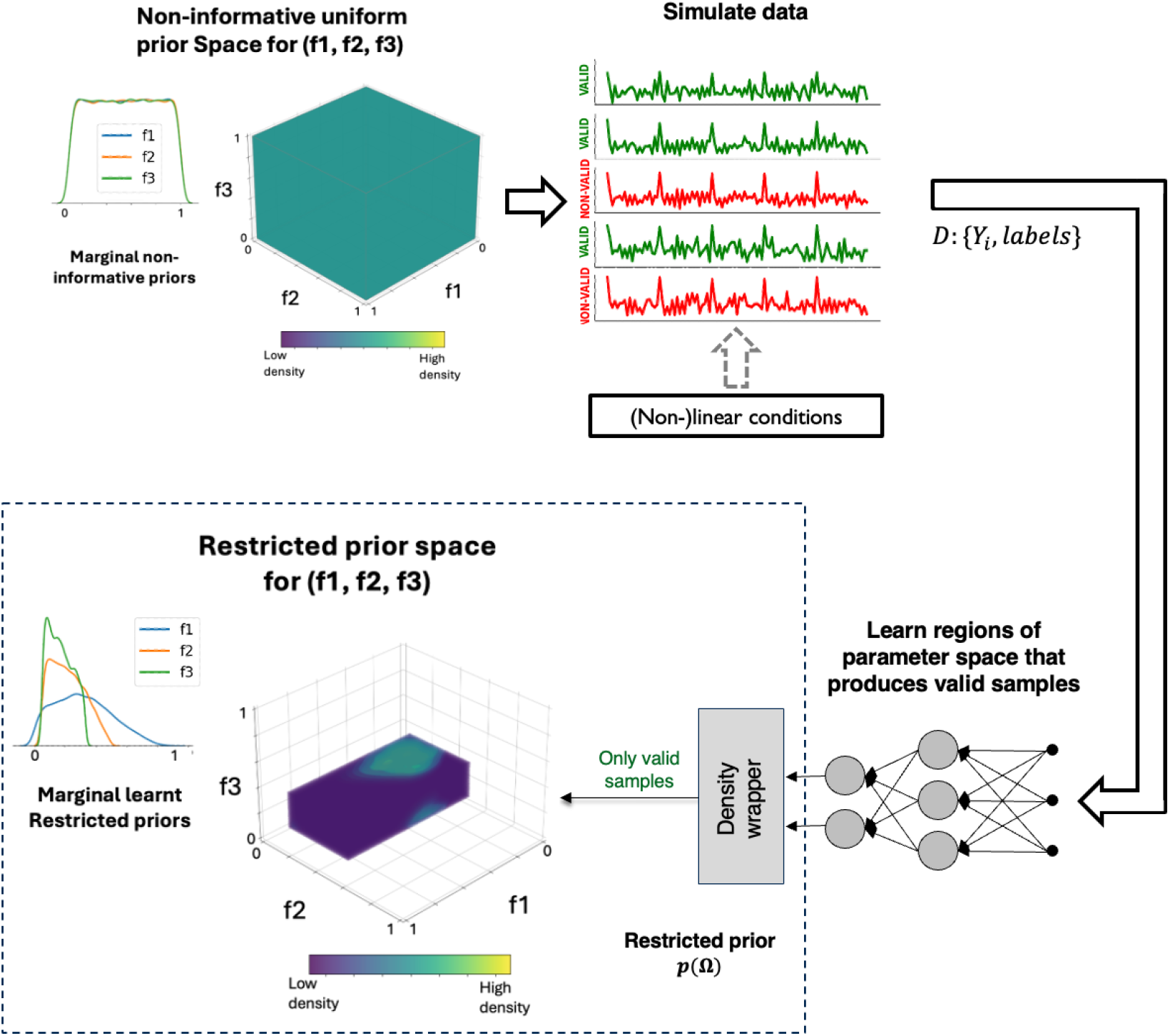
Restricted priors constraining the parameter space compared to non-informative uniform prior sampling. We simulated data *Y* using samples from the non-informative uniform priors and tagged these as “valid” or “non-valid” (according to the constraints we wanted to enforce) to create our training dataset *D*. A classifier was then trained to learn what is the signal stemming from “valid” parameter combinations and was used to produce samples only from the regions of the parameter space of interest i.e. a restricted prior. The figure shows an example of the learnt restricted priors used in this paper, which reflect the conditions and non-linear boundaries specified in Table 1 and target the high density areas; in this case, the marginal (top 2-D histograms) and the 3-D joint space of (*f*_1_, *f*_2_, *f*_3_) is shown. For a 3-fibre model, the sampling efficiency of prior samples is also increased with speed-ups of 105% compared to using a uniform sampling scheme and reject using the constraints as a post-sampling step.

**Figure S 3:**
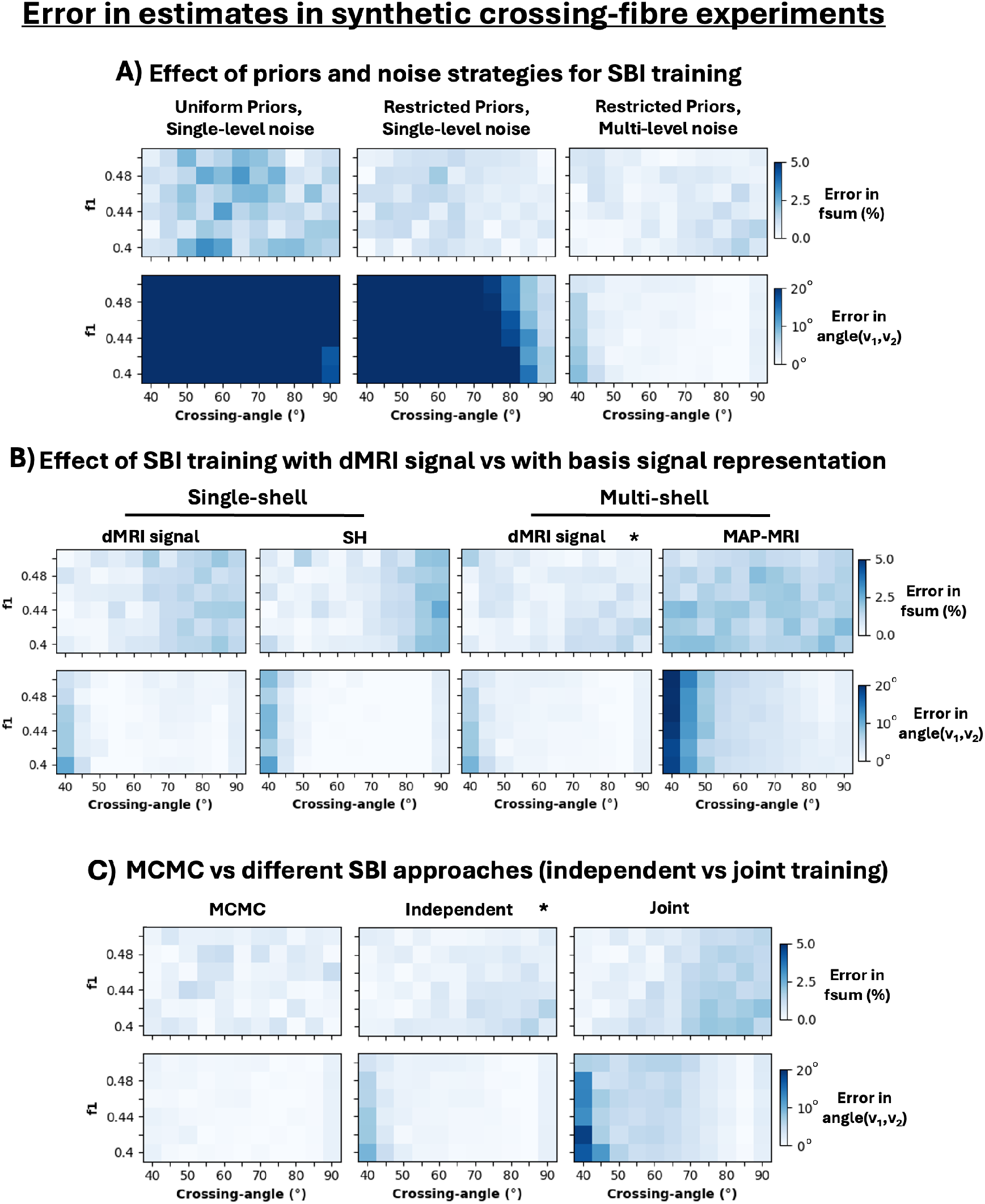
Evaluating accuracy of different SBI designs in synthetic fibre crossings (*N* = 2). dMRI was synthetically generated for patterns with two crossing fibres, varying the anisotropic volume fraction *f*_1_ and their crossing angle (rest of parameters were *d* = 0.0012 *mm*^2^*/s, f*_2_ = 0.2, *SNR* = 25 for all experiments). Median error over 100 noisy realisations, where the error is measured as 100 * (*groundtruth*_*i*_ − *estimate*_*i*_)*/groundtruth*_*i*_ [%], are shown for the same cases as in Fig. 3: A) Different prior definition and noise strategies for SBI training. B) SBI trained with raw dMRI signal (acquisition-specific) vs a basis set signal representation (acquisition-agnostic). C) Comparing MCMC fitting a model with *N* = 2 fibres against NPEs trained only with 2-fibres examples (Independent) or with a joint superset that contains both 1- and 2-fibres cases (Joint).

**Figure S 4:**
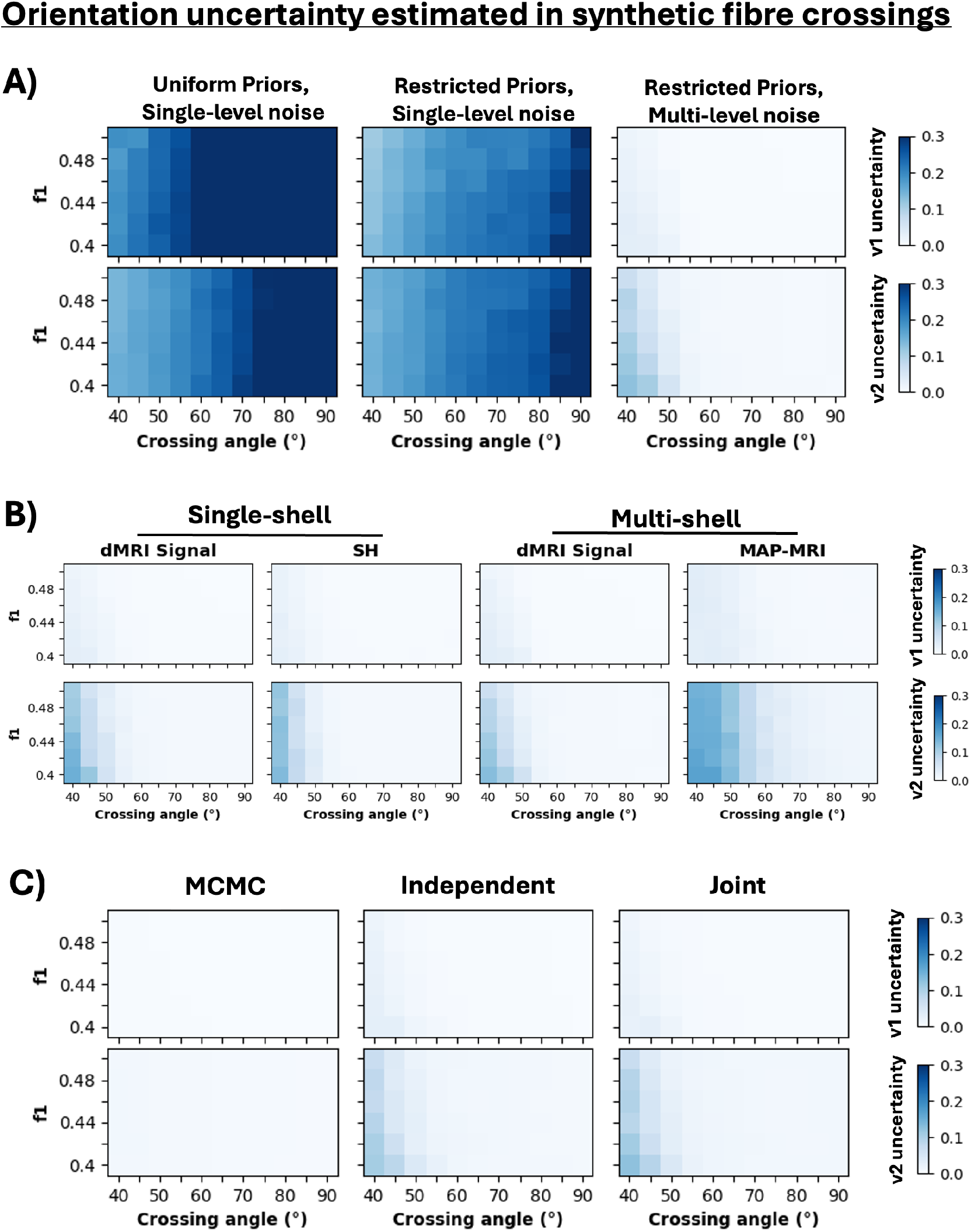
Evaluating precision of different SBI designs in mapping orientation uncertainty in synthetic fibre crossings (*N* = 2). dMRI was synthetically generated for patterns with two crossing fibres, varying the anisotropic volume fraction *f*_1_ and their crossing angle (rest of parameters were *d* = 0.0012 *mm*^2^*/s, f*_2_ = 0.2, *SNR* = 25 for all experiments). Uncertainty in **v**_1_ and **v**_2_ obtained from the width of the respective posteriors (median over 100 noisy realisations) are shown for the same cases as in Fig. 3 in the main text: A) Different prior definition and noise strategies for SBI training. B) SBI trained with raw dMRI signal (acquisition-specific) vs a basis set signal representation (acquisition-agnostic). C) Comparing MCMC fitting a model with *N* = 2 fibres against NPEs trained only with 2-fibres examples (Independent) or with a joint superset that contains both 1- and 2-fibres cases (Joint).

**Figure S 5:**
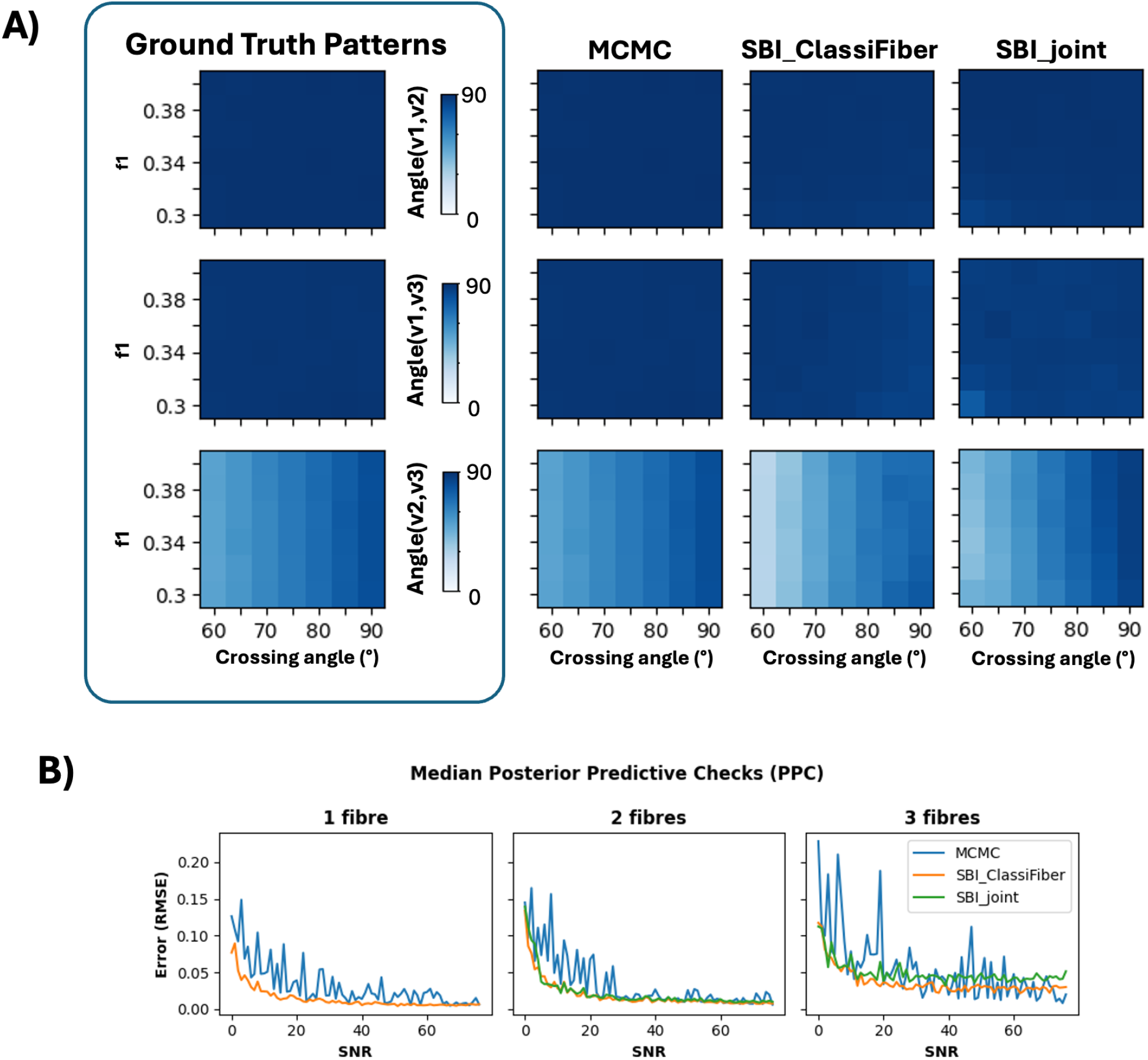
Evaluating performance of different SBI designs and MCMC in estimating crossing angles in synthetic 3-way fibre crossings (*N* = 3). A) Estimates from three approaches for synthetic data, where the first fibre orientation was perpendicular to the rest of compartments, and *f*_1_ and the crossing angle between the second and third fiber were varied. The rest of the parameters were kept to *d* = 0.0012, *f*_2_ = 0.25, *f*_3_ = 0.15, *SNR* = 25. B) Posterior Predictive Checks (PPC) as a function of SNR for synthetic experiments with different fiber complexities (*N* = 1, 2, 3) and parameter combinations. Each plot shows the relative error (median across 1,000 experiments) of the model predicted signal (using the mean posterior value for each parameter) with respect to the true signal.

**Figure S 6:**
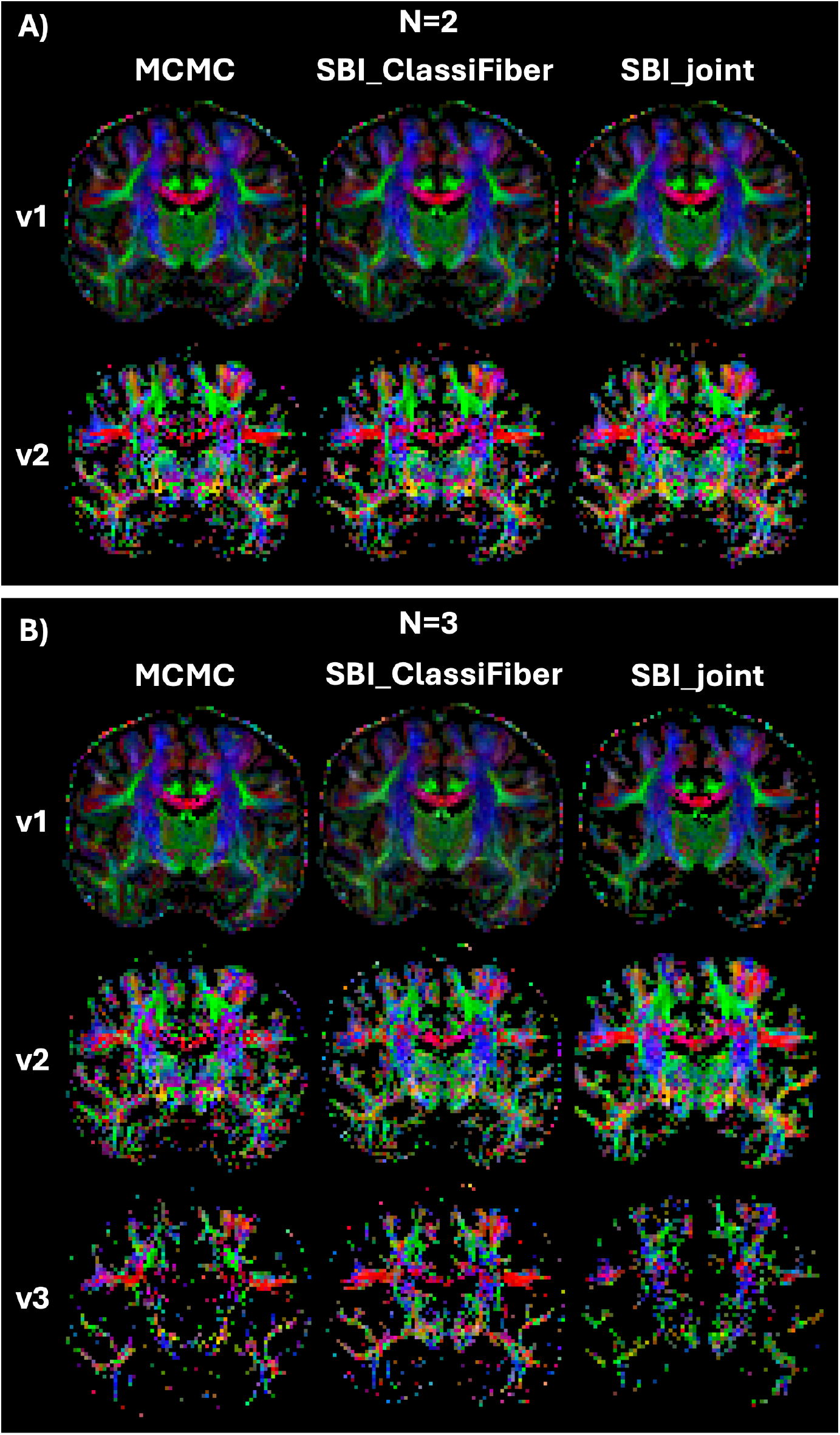
Maps of fiber orientation estimates (mean orientations of the posterior distributions) in in-vivo brain data obtained using MCMC, SBI_ClassiFiber and SBI_joint. A) Multi-shell 2-fibres model. B) Multi-shell 3-fibres model. In SBI, all samples have been corrected to avoid compartment swapping, as detailed in Fig. 7 in the main text.

**Figure S 7:**
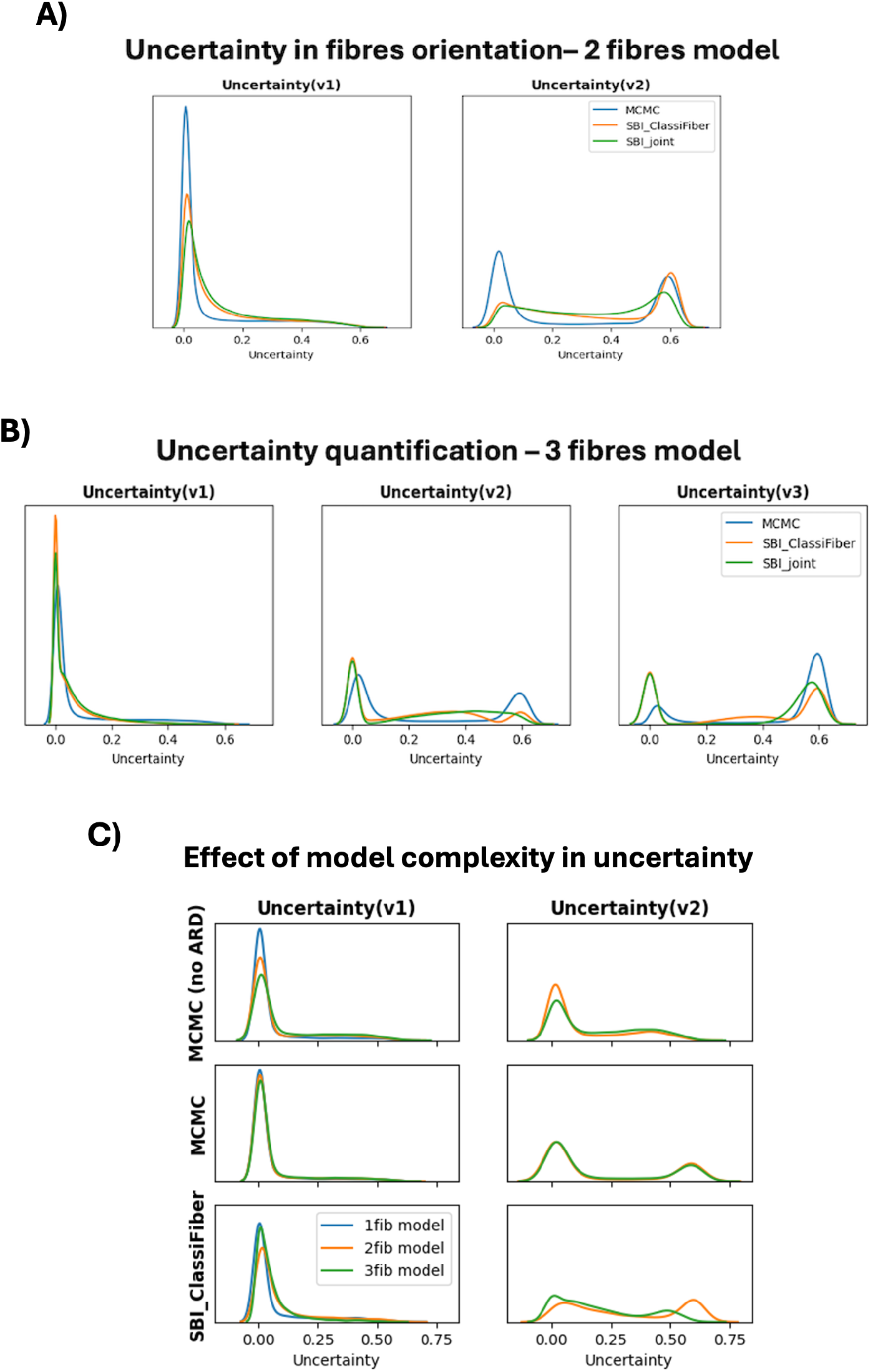
Comparison of uncertainty histograms returned by the different approaches. A) Uncertainty distribution in *v*_1_ and *v*_2_ mapped by the different approaches in the 2-fibres model. B) Uncertainty distribution in *v*_1_, *v*_2_ and *v*_3_ mapped by the different approaches in the 3-fibres model. C) Effect of increasing model complexity (models with *N* 1 to 3) in the uncertainty quantification of the first (*v*_1_) and second (*v*_2_) fibre orientation. Note the effect of the ARD priors in MCMC (first and second rows).

**Figure S 8:**
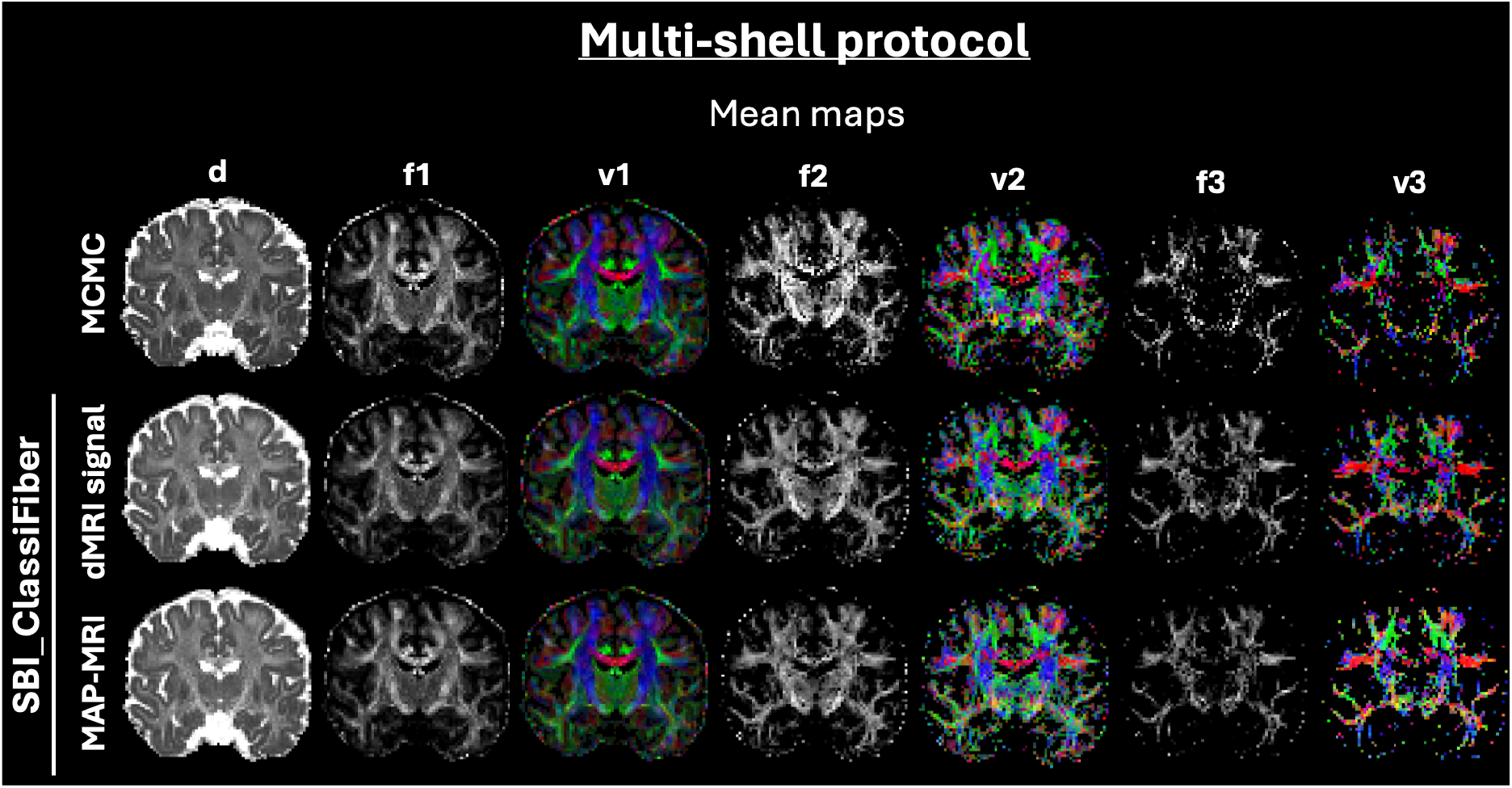
Maps of mean posterior estimates in in-vivo data from SBI trained against the raw dMRI signal (acquisition-specific) or a basis representation of the signal (acquisition=agnostic), for *N* = 3 compartments. MCMC estimates are shown for reference at the top row. Multi-shell data have been used and a MAP-MRI basis. NPEs are trained with 4 million samples and with the SBI_ClassiFiber architecture.

**Figure S 9:**
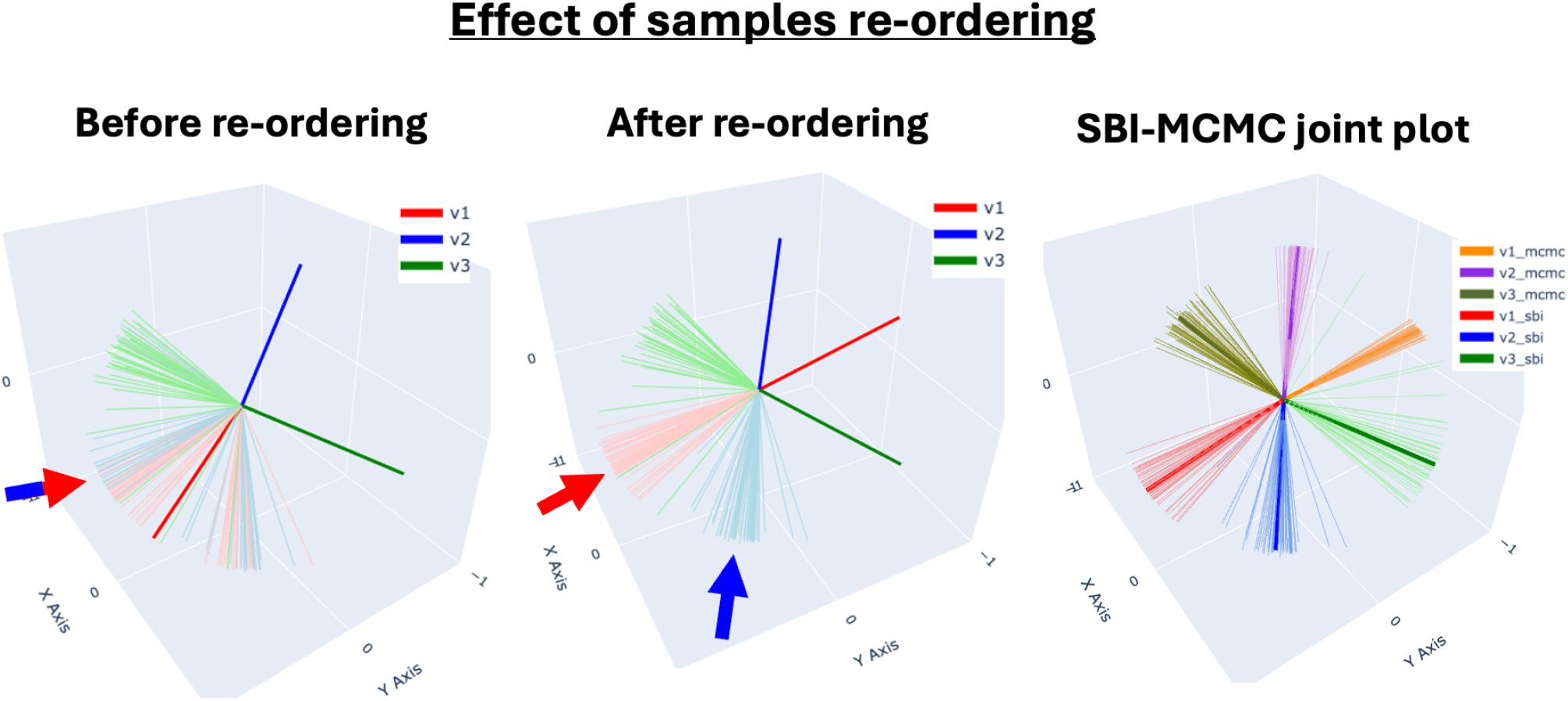
Effect of sample re-ordering in the joint distribution of fibre orientation samples. Example of a 3-way crossing obtained from in-vivo brain data in the Centrum Semiovale. Left: samples from the posterior provided by SBI. Middle: SBI samples re-grouped, based on similarity of orientations. Right: joint plot of the fibre orientation samples provided by MCMC and SBI after re-ordering. In all cases, samples represent the same posterior distribution, but the re-ordering of SBI samples helps in matching the mean vectors obtained by MCMC.

**Figure S 10:**
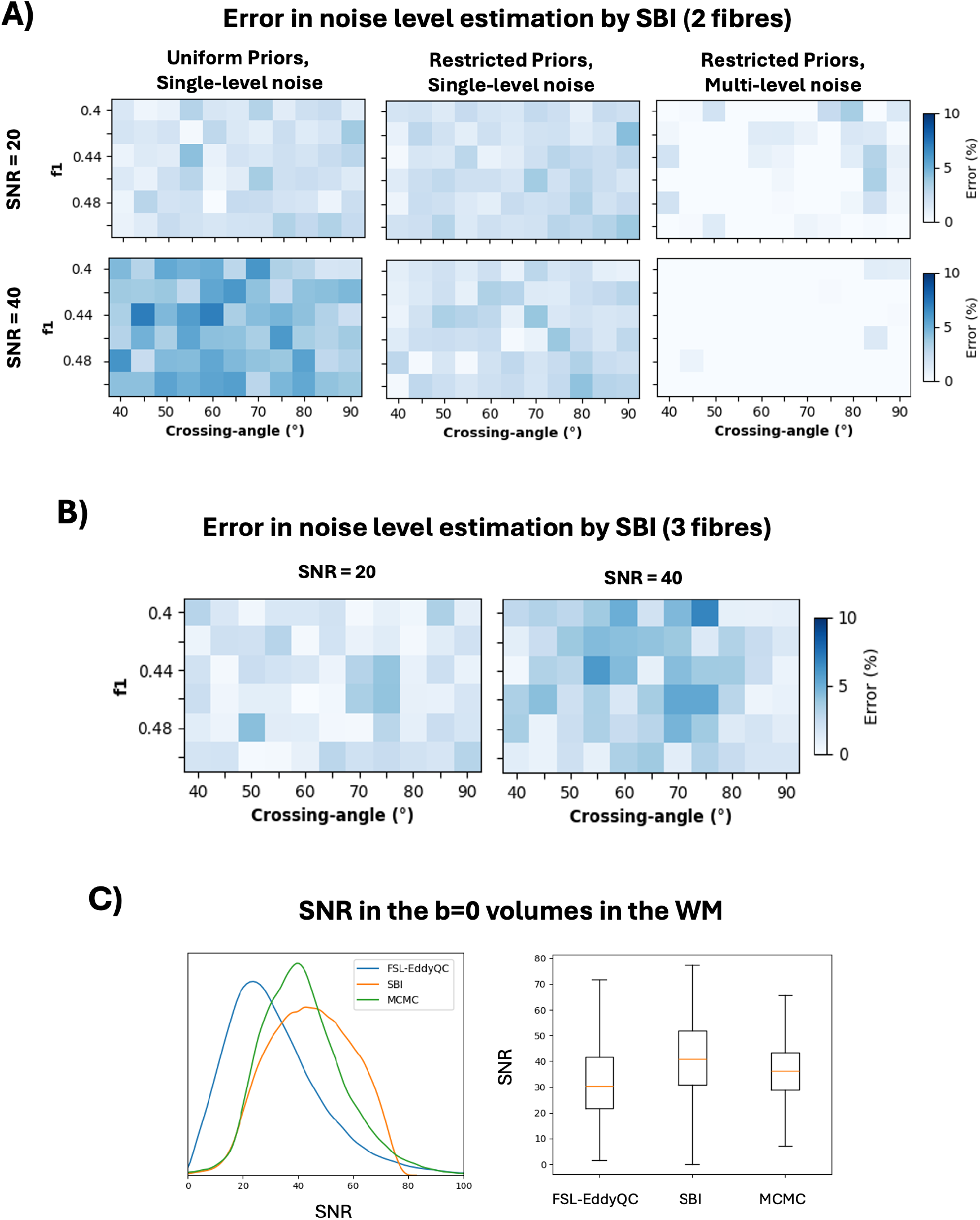
Noise estimation by SBI. A) Comparison of the error (in %) of the mean estimated SNR level by the different SBI designs, using the same models (i.e. *N* = 2) and synthetic data as in Fig.3 of main text at different noise levels. B) Error (in %) of the mean estimated SNR level using the same dataset as in panel A) but doing inference using the SBI_Classifiber with *N* = 3 (i.e. including model selection). C) Comparison of the distribution of mean SNR estimates in white matter obtained by MCMC and SBI, compared against SNR estimated directly from the b=0 s/mm^2^ volumes by FSL-EddyQC [82].

## References

[1] Dmitry S. Novikov. The present and the future of microstructure MRI: From a paradigm shift to normal science. Journal of Neuroscience Methods, 351, March 2021. ISSN 0165-0270. doi: 10.1016/j.jneumeth.2020.108947.

[2] Daniel C. Alexander, Tim B. Dyrby, Markus Nilsson, and Hui Zhang. Imaging brain microstructure with diffusion MRI: practicality and applications. NMR in Biomedicine, 32(4), April 2019. ISSN 0952-3480, 1099-1492. doi: 10.1002/nbm.3841.

[3] Saad Jbabdi, Stamatios N Sotiropoulos, Suzanne N Haber, David C Van Essen, and Timothy E Behrens. Measuring macroscopic brain connections in vivo. Nat Neurosci, 18(11):1546–1555, nov 2015. ISSN 1097-6256, 1546-1726. doi: 10.1038/nn.4134.

[4] Davood Karimi and Simon K. Warfield. Diffusion mri with machine learning. Imaging Neuroscience, 2:1–55, 11 2024. ISSN 2837-6056. doi: 10.1162/imag_a_00353.

[5] Derek K. Jones. Determining and visualizing uncertainty in estimates of fiber orientation from diffusion tensor MRI. Magn. Reson. Med., 49(1):7–12, jan 2003. ISSN 0740-3194, 1522-2594. doi: 10.1002/mrm.10331.

[6] Daniel C. Alexander. A general framework for experiment design in diffusion MRI and its application in measuring direct tissue-microstructure features. Magn. Reson. Med., 60(2):439–448, aug 2008. ISSN 07403194, 15222594. doi: 10.1002/mrm.21646.

[7] T.E.J. Behrens, H. Johansen Berg, S. Jbabdi, M.F.S. Rushworth, and M.W. Woolrich. Probabilistic diffusion tractography with multiple fibre orientations: What can we gain? NeuroImage, 34(1):144–155, jan 2007. ISSN 10538119. doi: 10.1016/j.neuroimage.2006.09.018.

[8] Enrico Kaden, Thomas R. Knösche, and Alfred Anwander. Parametric spherical deconvolution: Inferring anatomical connectivity using diffusion MR imaging. NeuroImage, 37(2):474–488, aug 2007. ISSN 10538119. doi: 10.1016/j.neuroimage.2007.05.012.

[9] Anastasia Yendiki, Patricia Panneck, Priti Srinivasan, Allison Stevens, Lilla Zöllei, Jean Augustinack, Ruopeng Wang, David Salat, Stefan Ehrlich, Tim Behrens, Saad Jbabdi, Randy Gollub, and Bruce Fischl. Automated Probabilistic Reconstruction of White-Matter Pathways in Health and Disease Using an Atlas of the Underlying Anatomy. Frontiers in Neuroinformatics, 5:23, October 2011. doi: 10.3389/fninf.2011.00023.

[10] G. Girard, D. B. Aydogan, F. Dell’Acqua, A. Leemans, M. Descoteaux, and S. N. Sotiropoulos. 14. probabilistic tractography. In F. Dell’Acqua, A. Leemans, and M. Descoteaux, editors, Handbook of Diffusion MRI Tractography. Elsevier, 2024.

[11] Sinisa Pajevic and Peter J Basser. Parametric and non-parametric statistical analysis of DTMRI data. Journal of Magnetic Resonance, 161(1):1–14, March 2003. ISSN 10907807. doi: 10.1016/S1090-7807(02)00178-7.

[12] Derek K. Jones. Tractography Gone Wild: Probabilistic Fibre Tracking Using the Wild Bootstrap With Diffusion Tensor MRI. IEEE Transactions on Medical Imaging, 27(9):1268–1274, September 2008. ISSN 1558-254X. doi: 10.1109/TMI.2008.922191.

[13] Adam S. Bernstein, Nan-kuei Chen, and Theodore P. Trouard. Bootstrap analysis of diffusion tensor and mean apparent propagator parameters derived from multiband diffusion MRI. Magn Reson Med, 82(5):1796–1803, November 2019. ISSN 0740-3194. doi: 10.1002/mrm.27833.

[14] Xuan Gu, Anders Eklund, Evren Özarslan, and Hans Knutsson. Using the Wild Bootstrap to Quantify Uncertainty in Mean Apparent Propagator MRI. Frontiers in Neuroinformatics, 13, 2019. ISSN 1662-5196. doi: 10.3389/fninf.2019.00043.

[15] Lipeng Ning, Filip Szczepankiewicz, Markus Nilsson, Yogesh Rathi, and Carl-Fredrik Westin. Probing tissue microstructure by diffusion skewness tensor imaging. Sci Rep, 11(1):135, December 2021. ISSN 2045-2322. doi: 10.1038/s41598-020-79748-3.

[16] Brandon Whitcher, David S. Tuch, Jonathan J. Wisco, A. Gregory Sorensen, and Liqun Wang. Using the wild bootstrap to quantify uncertainty in diffusion tensor imaging. Hum Brain Mapp, 29 (3):346–362, March 2008. ISSN 1065-9471. doi: 10.1002/hbm.20395.

[17] Göran Kauermann, Gerda Claeskens, and J. D. Opsomer. Bootstrapping for Penalized Spline Regression. Journal of Computational and Graphical Statistics, 18(1):126–146, 2009. ISSN 1061-8600. URL https://www.jstor.org/stable/25703557.

[18] Michael A. Chappell, Adrian R. Groves, Brandon Whitcher, and Mark W. Woolrich. Variational Bayesian Inference for a Nonlinear Forward Model. IEEE Transactions on Signal Processing, 57(1):223–236, January 2009. ISSN 1941-0476. doi: 10.1109/TSP.2008.2005752.

[19] Enrico Kaden, Alfred Anwander, and Thomas R. Knösche. Variational inference of the fiber orientation density using diffusion MR imaging. NeuroImage, 42(4):1366–1380, October 2008. ISSN 10538119. doi: 10.1016/j.neuroimage.2008.06.004.

[20] David M. Blei, Alp Kucukelbir, and Jon D. McAuliffe. Variational Inference: A Review for Statisticians. Journal of the American Statistical Association, (518):859–877, April 2017. ISSN 0162-1459, 1537-274X. doi: 10.1080/01621459.2017.1285773. 1601.00670.

[21] T.E.J. Behrens, M.W. Woolrich, M. Jenkinson, H. Johansen-Berg, R.G. Nunes, S. Clare, P.M. Matthews, J.M. Brady, and S.M. Smith. Characterization and propagation of uncertainty in diffusion-weighted MR imaging. Magn. Reson. Med., 50(5):1077–1088, November 2003. ISSN 0740-3194, 1522-2594. doi: 10.1002/mrm.10609.

[22] Enrico Kaden and Frithjof Kruggel. Nonparametric Bayesian inference of the fiber orientation distribution from diffusion-weighted MR images. Medical Image Analysis, 16(4):876–888, May 2012. ISSN 13618415. doi: 10.1016/j.media.2012.01.004.

[23] S. N. Sotiropoulos, S. Jbabdi, J. L. Andersson, M. W. Woolrich, K. Ugurbil, and T. E. J. Behrens. RubiX: Combining Spatial Resolutions for Bayesian Inference of Crossing Fibers in Diffusion MRI. IEEE Trans. Med. Imaging, 32(6):969–982, June 2013. ISSN 0278-0062, 1558-254X. doi: 10.1109/TMI.2012.2231873.

[24] Moises Hernandez-Fernandez, Istvan Reguly, Saad Jbabdi, Mike Giles, Stephen Smith, and Stamatios N. Sotiropoulos. Using GPUs to accelerate computational diffusion MRI: From microstructure estimation to tractography and connectomes. NeuroImage, 188:598–615, March 2019. ISSN 10538119. doi: 10.1016/j.neuroimage.2018.12.015.

[25] Jose Pedro Manzano Patron. On noise, uncertainty and inference for computational diffusion MRI. PhD Thesis, University of Nottingham, 2023. URL http://eprints.nottingham.ac.uk/74189/.

[26] R. L. Harms, F. J. Fritz, S. Schoenmakers, and A. Roebroeck. Fast and robust quantification of uncertainty in non-linear diffusion MRI models. NeuroImage, 285:120496, January 2024. ISSN 1053-8119. doi: 10.1016/j.neuroimage.2023.120496.

[27] Kyle Cranmer, Johann Brehmer, and Gilles Louppe. The frontier of simulation-based inference. Proc Natl Acad Sci USA, 117(48):30055–30062, December 2020. ISSN 0027-8424, 1091-6490. doi: 10.1073/pnas.1912789117.

[28] Stefan T Radev, Ulf K Mertens, Andreas Voss, Lynton Ardizzone, and Ullrich K”othe. Bayesflow: Learning complex stochastic models with invertible neural networks. IEEE transactions on neural networks and learning systems, 33(4):1452–1466, 2020.

[29] Joeri Hermans, Volodimir Begy, and Gilles Louppe. Likelihood-free mcmc with amortized approximate ratio estimators. In International conference on machine learning, pages 4239–4248. PMLR, 2020.

[30] Saad Jbabdi, Stamatios N. Sotiropoulos, Alexander M. Savio, Manuel Graña, and Timothy E. J. Behrens. Model-based analysis of multishell diffusion MR data for tractography: How to get over fitting problems. Magnetic Resonance in Medicine, 68(6):1846–1855, 2012. ISSN 1522-2594. doi: 10.1002/mrm.24204.

[31] Moisés Hernández, Ginés D. Guerrero, José M. Cecilia, José M. García, Alberto Inuggi, Saad Jbabdi, Timothy E. J. Behrens, and Stamatios N. Sotiropoulos. Accelerating Fibre Orientation Estimation from Diffusion Weighted Magnetic Resonance Imaging Using GPUs. PLOS ONE, 8(4):e61892, 2013. ISSN 1932-6203. doi: 10.1371/journal.pone.0061892.

[32] S. Tavaré, D. J. Balding, R. C. Griffiths, and P. Donnelly. Inferring coalescence times from DNA sequence data. Genetics, 145(2):505–518, February 1997. ISSN 0016-6731. doi: 10.1093/genetics/145.2.505.

[33] Mark A. Beaumont. Approximate Bayesian Computation in Evolution and Ecology. Annual Review of Ecology, Evolution, and Systematics, 41(Volume 41, 2010):379–406, December 2010. ISSN 1543-592X, 1545-2069. doi: 10.1146/annurev-ecolsys-102209-144621.

[34] S. A. Sisson, Y. Fan, and M. A. Beaumont. Overview of Approximate Bayesian Computation. 1802.09720 [stat], February 2018. URL http://arxiv.org/abs/1802.09720.

[35] Mark A Beaumont, Wenyang Zhang, and David J Balding. Approximate Bayesian Computation in Population Genetics. Genetics, 162(4):2025–2035, December 2002. ISSN 1943-2631. doi: 10.1093/genetics/162.4.2025.

[36] Michael G. B. Blum and Olivier François. Non-linear regression models for Approximate Bayesian Computation. Stat Comput, 20(1):63–73, March 2009. ISSN 1573-1375. doi: 10.1007/s11222-009-9116-0.

[37] Malik Magdon-Ismail and Amir Atiya. Neural Networks for Density Estimation. In Advances in Neural Information Processing Systems, volume 11. MIT Press, 1998. URL https://proceedings.neurips.cc/paper_files/paper/1998/hash/9327969053c0068dd9e07c529866b94d-Abstract.html.

[38] George Papamakarios and Iain Murray. Fast $\epsilon$-Free Inference of Simulation Models with Bayesian Conditional Density Estimation. In Advances in Neural Information Processing Systems. Curran Associates, Inc., 2016. URL https://proceedings.neurips.cc/paper_files/paper/2016/hash/6aca97005c68f1206823815f66102863-Abstract.html.

[39] Jan-Matthis Lueckmann, Pedro J Goncalves, Giacomo Bassetto, Kaan Öcal, Marcel Nonnenmacher, and Jakob H Macke. Flexible statistical inference for mechanistic models of neural dynamics. In Advances in Neural Information Processing Systems, volume 30. Curran Associates, Inc., 2017. URL https://proceedings.neurips.cc/paper/2017/hash/addfa9b7e234254d26e9c7f2af1005cb-Abstract.html.

[40] George Papamakarios, Theo Pavlakou, and Iain Murray. Masked autoregressive flow for density estimation. In I. Guyon, U. Von Luxburg, S. Bengio, H. Wallach, R. Fergus, S. Vishwanathan, and R. Garnett, editors, Advances in Neural Information Processing Systems, volume 30. Curran Associates, Inc., 2017. URL https://proceedings.neurips.cc/paper_files/paper/2017/file/6c1da886822c67822bcf3679d04369fa-Paper.pdf.

[41] David Greenberg, Marcel Nonnenmacher, and Jakob Macke. Automatic Posterior Transformation for Likelihood-Free Inference. In Proceedings of the 36th International Conference on Machine Learning, pages 2404–2414. PMLR, May 2019. URL https://proceedings.mlr.press/v97/greenberg19a.html. ISSN: 2640-3498.

[42] George Papamakarios, David Sterratt, and Iain Murray. Sequential neural likelihood: Fast likelihood-free inference with autoregressive flows. In Kamalika Chaudhuri and Masashi Sugiyama, editors, Proceedings of the Twenty-Second International Conference on Artificial Intelligence and Statistics, volume 89 of Proceedings of Machine Learning Research, pages 837–848. PMLR, 16–18 Apr 2019. URL https://proceedings.mlr.press/v89/papamakarios19a.html.

[43] Conor Durkan, George Papamakarios, and Iain Murray. Sequential Neural Methods for Likelihood-free Inference, November 2018. URL http://arxiv.org/abs/1811.08723.

[44] Louis Sharrock, Jack Simons, Song Liu, and Mark Beaumont. Sequential neural score estimation: Likelihood-free inference with conditional score based diffusion models. arXiv preprint, 2022. doi: 2210.04872.

[45] Tomas Geffner, George Papamakarios, and Andriy Mnih. Compositional score modeling for simulation-based inference. In International Conference on Machine Learning, pages 11098–11116. PMLR, 2023.

[46] Jonas Wildberger, Maximilian Dax, Simon Buchholz, Stephen Green, Jakob H Macke, and Bernhard Schölkopf. Flow matching for scalable simulation-based inference. Advances in Neural Information Processing Systems, 36, 2024.

[47] Manuel Gloeckler, Michael Deistler, Christian Dietrich Weilbach, Frank Wood, and Jakob H Macke. All-in-one simulation-based inference. In Forty-first International Conference on Machine Learning, 2024.

[48] Maëliss Jallais, Pedro L. C. Rodrigues, Alexandre Gramfort, and Demian Wassermann. Inverting brain grey matter models with likelihood-free inference: a tool for trustable cytoarchitecture measurements. Melba, 1(IPMI 2021):1–28, May 2022. ISSN 2766-905X. doi: 10.59275/j.melba.2022-a964.

[49] Marco Palombo, Andrada Ianus, Michele Guerreri, Daniel Nunes, Daniel C. Alexander, Noam Shemesh, and Hui Zhang. SANDI: A compartment-based model for non-invasive apparent soma and neurite imaging by diffusion MRI. Neuroimage, 215:116835, July 2020. ISSN 1095-9572. doi: 10.1016/j.neuroimage.2020.116835.

[50] Maeliss Jallais and Marco Palombo. mu-GUIDE: a framework for quantitative imaging via generalized uncertainty-driven inference using deep learning. eLife, 13, October 2024. doi: 10.7554/eLife.101069.2.

[51] Maximilian F. Eggl and Silvia De Santis. More with less: Simulation-based inference enables accurate diffusion-weighted MRI with minimal acquisition time, November 2024.

[52] Hazhar Sufi Karimi, Arghya Pal, Lipeng Ning, and Yogesh Rathi. Likelihood-free posterior estimation and uncertainty quantification for diffusion MRI models. Imaging Neuroscience, 2:1–22, February 2024. ISSN 2837-6056. doi: 10.1162/imag_a_00088.

[53] Dmitry S. Novikov, Jelle Veraart, Ileana O. Jelescu, and Els Fieremans. Rotationally-invariant mapping of scalar and orientational metrics of neuronal microstructure with diffusion MRI. NeuroImage, 174:518–538, July 2018. ISSN 1053-8119. doi: 10.1016/j.neuroimage.2018.03.006.

[54] William Consagra, Lipeng Ning, and Yogesh Rathi. A Deep Learning Approach to Multi-Fiber Parameter Estimation and Uncertainty Quantification in Diffusion MRI, May 2024.

[55] Jose Pedro Manzano-Patron, Theodore Kypraios, and Stamatios N. Sotiropoulos. Amortised inference in diffusion mri biophysical models using artificial neural networks and simulation-based frameworks. In ISMRM (Proc. Intl. Soc. Mag. Reson. Med. 30), 2022. doi: 10.58530/2022/1644.

[56] George Papamakarios, Eric Nalisnick, Danilo Jimenez Rezende, Shakir Mohamed, and Balaji Lakshminarayanan. Normalizing Flows for Probabilistic Modeling and Inference. Journal of Machine Learning Research, 22(57):1–64, 2021. ISSN 1533-7928. URL http://jmlr.org/papers/v22/19-1028.html.

[57] Jan-Matthis Lueckmann, Jan Boelts, David Greenberg, Pedro Goncalves, and Jakob Macke. Benchmarking Simulation-Based Inference. In Proceedings of The 24th International Conference on Artificial Intelligence and Statistics, pages 343–351. PMLR, March 2021. URL https://proceedings.mlr.press/v130/lueckmann21a.html.

[58] Tim Dockhorn, James A. Ritchie, Yaoliang Yu, and Iain Murray. Density Deconvolution with Normalizing Flows. 2006.09396 [cs, stat], July 2020. URL http://arxiv.org/abs/2006.09396.

[59] Conor Durkan, Artur Bekasov, Iain Murray, and George Papamakarios. Neural Spline Flows. In Advances in Neural Information Processing Systems, volume 32. Curran Associates, Inc., 2019. URL https://proceedings.neurips.cc/paper_files/paper/2019/hash/7ac71d433f282034e088473244df8c02-Abstract.html.

[60] Alvaro Tejero-Cantero, Jan Boelts, Michael Deistler, Jan-Matthis Lueckmann, Conor Durkan, Pedro Gonçalves, David Greenberg, and Jakob Macke. sbi: A toolkit for simulation-based inference. JOSS, 5(52):2505, August 2020. ISSN 2475-9066. doi: 10.21105/joss.02505.

[61] Michael Deistler, Jakob H. Macke, and Pedro J. Gonçalves. Energy-efficient network activity from disparate circuit parameters. Proceedings of the National Academy of Sciences, 119(44):e2207632119, November 2022. doi: 10.1073/pnas.2207632119.

[62] Michael Deistler, Pedro J. Goncalves, and Jakob H. Macke. Truncated proposals for scalable and hassle-free simulation-based inference. In Thirty-Sixth Conference on Neural Information Processing Systems, 2022. URL https://openreview.net/forum?id=QW98XBAqNRa.

[63] Yoshitaka Masutani. Noise Level Matching Improves Robustness of Diffusion Mri Parameter Inference by Synthetic Q-Space Learning. In 2019 IEEE 16th International Symposium on Biomedical Imaging (ISBI 2019), pages 139–142, April 2019. doi: 10.1109/ISBI.2019.8759161. ISSN: 1945-8452.

[64] Cornelius Schröder and Jakob H. Macke. Simultaneous identification of models and parameters of scientific simulators, May 2024.

[65] Jose Pedro Manzano Patron, Steen Moeller, Jesper L.R. Andersson, Kamil Ugurbil, Essa Yacoub, and Stamatios N. Sotiropoulos. Denoising diffusion MRI: Considerations and implications for analysis. Imaging Neuroscience, 2:1–29, January 2024. ISSN 2837-6056. doi: 10.1162/imag_a_00060.

[66] Karla L Miller, Fidel Alfaro-Almagro, Neal K Bangerter, David L Thomas, Essa Yacoub, Junqian Xu, Andreas J Bartsch, Saad Jbabdi, Stamatios N Sotiropoulos, Jesper LR Andersson, Ludovica Griffanti, Gwenaëlle Douaud, Thomas W Okell, Peter Weale, Iulius Dragonu, Steve Garratt, Sarah Hudson, Rory Collins, Mark Jenkinson, Paul M Matthews, and Stephen M Smith. Multimodal population brain imaging in the UK Biobank prospective epidemiological study. Nat Neurosci, 19 (11):1523–1536, November 2016. ISSN 1097-6256, 1546-1726. doi: 10.1038/nn.4393.

[67] Ben Jeurissen, Alexander Leemans, Jacques-Donald Tournier, Derek K. Jones, and Jan Sijbers. Investigating the prevalence of complex fiber configurations in white matter tissue with diffusion magnetic resonance imaging: Prevalence of Multifiber Voxels in WM. Hum. Brain Mapp, 34(11):2747–2766, November 2013. ISSN 10659471. doi: 10.1002/hbm.22099.

[68] Maxime Descoteaux, Elaine Angelino, Shaun Fitzgibbons, and Rachid Deriche. Regularized, fast, and robust analytical Q-ball imaging. Magnetic Resonance in Medicine, 58(3):497–510, 2007. ISSN 1522-2594. doi: 10.1002/mrm.21277.

[69] Evren Özarslan, Cheng Guan Koay, Timothy M. Shepherd, Michal E. Komlosh, M. Okan Irfanoglu, Carlo Pierpaoli, and Peter J. Basser. Mean apparent propagator (MAP) MRI: a novel diffusion imaging method for mapping tissue microstructure. Neuroimage, 78:16–32, September 2013. ISSN 1095-9572. doi: 10.1016/j.neuroimage.2013.04.016.

[70] Eleftherios Garyfallidis, Matthew Brett, Bagrat Amirbekian, Ariel Rokem, Stefan van der Walt, Maxime Descoteaux, Ian Nimmo-Smith, and Dipy Contributors. Dipy, a library for the analysis of diffusion MRI data. Front. Neuroinform., 8, February 2014. ISSN 1662-5196. doi: 10.3389/fninf.2014.00008.

[71] Shaun Warrington, Katherine L. Bryant, Alexandr A. Khrapitchev, Jerome Sallet, Marina Charquero-Ballester, Gwenaëlle Douaud, Saad Jbabdi, Rogier B. Mars, and Stamatios N. Sotiropoulos. XTRACT - Standardised protocols for automated tractography in the human and macaque brain. NeuroImage, 217:116923, August 2020. ISSN 10538119. doi: 10.1016/j.neuroimage.2020.116923.

[72] Meysam Hashemi, Anirudh N. Vattikonda, Jayant Jha, Viktor Sip, Marmaduke M. Woodman, Fabrice Bartolomei, and Viktor K. Jirsa. Amortized Bayesian inference on generative dynamical network models of epilepsy using deep neural density estimators. Neural Networks, 163:178–194, June 2023. ISSN 0893-6080. doi: 10.1016/j.neunet.2023.03.040.

[73] Jonas Rothfuss, Fabio Ferreira, Simon Boehm, Simon Walther, Maxim Ulrich, Tamim Asfour, and Andreas Krause. Noise Regularization for Conditional Density Estimation, February 2020. URL http://arxiv.org/abs/1907.08982.

[74] Joeri Hermans, Arnaud Delaunoy, François Rozet, Antoine Wehenkel, and Gilles Louppe. Averting A Crisis In Simulation-Based Inference, October 2021.

[75] Stefan T Radev, Marco D’Alessandro, Ulf K Mertens, Andreas Voss, Ullrich K”othe, and Paul-Christian B”urkner. Amortized bayesian model comparison with evidential deep learning. IEEE Transactions on Neural Networks and Learning Systems, 34(8):4903–4917, 2021.

[76] David P Wipf and Srikantan S Nagarajan. A New View of Automatic Relevance Determination. page 8, 2007.

[77] Mark W. Woolrich, Saad Jbabdi, Brian Patenaude, Michael Chappell, Salima Makni, Timothy Behrens, Christian Beckmann, Mark Jenkinson, and Stephen M. Smith. Bayesian analysis of neuroimaging data in FSL. NeuroImage, 45(1):S173–S186, March 2009. ISSN 10538119. doi: 10.1016/j.neuroimage.2008.10.055.

[78] Thomas Minka. Divergence measures and message passing. 2005.

[79] Conor Durkan, Iain Murray, and George Papamakarios. On Contrastive Learning for Likelihood-free Inference. In Proceedings of the 37th International Conference on Machine Learning, pages 2771– 2781. PMLR, November 2020. URL https://proceedings.mlr.press/v119/durkan20a.html. ISSN: 2640-3498.

[80] Laurence Illing Midgley, Vincent Stimper, Gregor N. C. Simm, Bernhard Schölkopf, and José Miguel Hernández-Lobato. Flow Annealed Importance Sampling Bootstrap, March 2023.

[81] Maximilian Dax, Stephen R. Green, Jonathan Gair, Michael Pürrer, Jonas Wildberger, Jakob H. Macke, Alessandra Buonanno, and Bernhard Schölkopf. Neural Importance Sampling for Rapid and Reliable Gravitational-Wave Inference. Phys. Rev. Lett., 130(17):171403, April 2023. doi: 10.1103/PhysRevLett.130.171403.

[82] Matteo Bastiani, Michiel Cottaar, Sean P. Fitzgibbon, Sana Suri, Fidel Alfaro-Almagro, Stamatios N. Sotiropoulos, Saad Jbabdi, and Jesper L.R. Andersson. Automated quality control for within and between studies diffusion MRI data using a non-parametric framework for movement and distortion correction. NeuroImage, 184:801–812, January 2019. ISSN 10538119. doi: 10.1016/j.neuroimage.2018.09.073. Citation Key Alias: 2019bastianiAutomatedQualityControl.

[83] Poornima Ramesh, Jan-Matthis Lueckmann, Jan Boelts, Álvaro Tejero-Cantero, David S. Greenberg, Pedro J. Gonçalves, and Jakob H. Macke. GATSBI: Generative Adversarial Training for Simulation-Based Inference. In International Conference on Learning Representations, 2022. URL https://openreview.net/pdf?id=kR1hC6j48Tp.

[84] William Consagra, Lipeng Ning, and Yogesh Rathi. Neural orientation distribution fields for estimation and uncertainty quantification in diffusion MRI. Medical Image Analysis, 93:103105, April 2024. ISSN 1361-8415. doi: 10.1016/j.media.2024.103105.

[85] Murat Demirtaş, Joshua B. Burt, Markus Helmer, Jie Lisa Ji, Brendan D. Adkinson, Matthew F. Glasser, David C. Van Essen, Stamatios N. Sotiropoulos, Alan Anticevic, and John D. Murray. Hierarchical Heterogeneity across Human Cortex Shapes Large-Scale Neural Dynamics. Neuron, 101(6):1181–1194.e13, March 2019. ISSN 0896-6273. doi: 10.1016/j.neuron.2019.01.017.

[86] Viktor Jirsa, Huifang Wang, Paul Triebkorn, Meysam Hashemi, Jayant Jha, Jorge Gonzalez-Martinez, Maxime Guye, Julia Makhalova, and Fabrice Bartolomei. Personalised virtual brain models in epilepsy. The Lancet Neurology, 22(5):443–454, May 2023. ISSN 1474-4422. doi: 10.1016/S1474-4422(23)00008-X.

[87] Harith Akram, Stamatios N. Sotiropoulos, Saad Jbabdi, Dejan Georgiev, Philipp Mahlknecht, Jonathan Hyam, Thomas Foltynie, Patricia Limousin, Enrico De Vita, Marjan Jahanshahi, Marwan Hariz, John Ashburner, Tim Behrens, and Ludvic Zrinzo. Subthalamic deep brain stimulation sweet spots and hyperdirect cortical connectivity in Parkinson’s disease. NeuroImage, 158:332–345, September 2017. ISSN 1053-8119. doi: 10.1016/j.neuroimage.2017.07.012.

[88] Raymond Salvador, Alonso Peña, David K. Menon, T. Adrian Carpenter, John D. Pickard, and Ed T. Bullmore. Formal characterization and extension of the linearized diffusion tensor model: Linearized Diffusion Tensor Model. Hum. Brain Mapp., 24(2):144–155, February 2005. ISSN 10659471. doi: 10.1002/hbm.20076.

